# Tamoxifen and Doxorubicin Synergistically Target Estrogen Receptor β-Mutant p53-p73 Axis, a Therapeutically Actionable Vulnerability in Triple Negative Breast Cancer

**DOI:** 10.64898/2026.10.21.760572

**Authors:** Chetan C Oturkar, Junhoung Park J, Iqbal Ijaz, Christina Adams, Utpal K Mukhopadhyay, Melissa Dolan, Alexander Caradori, Manasori Oshi, Yali Zhang, Jianmin Wang, Brent Boleslav, Donald Mager, Abhilash Nair, Tagari Samanta, Kwang Hwa Jung, Seung Yeop Han, Sukjin Yang, Abha Tiwari, Richa Mishra, John Ebos, Kazuaki Takabe, Kris Attwood, Benny A Kaipparettu, Gokul M Das

## Abstract

Current therapeutic regimens for triple negative breast cancer (TNBC) are not very effective in majority of patients and have serious adverse effects. Tamoxifen (Tam), a selective estrogen receptor modulator (SERM), routinely used to treat estrogen receptor alpha (ERα)-positive breast cancer, has traditionally not been considered for TNBC therapy because it lacks expression of ERα, the canonical target of Tam. Challenging this conventional clinical paradigm, we have demonstrated, using genetic and pharmacological approaches, that Tam synergizes with DNA-damaging agent doxorubicin (Doxo) to increase cell death by both apoptosis and ferroptosis leading to growth inhibition of TNBC CDXs and PDXs expressing mutant p53. Mechanistically, these responses are dependent on ERβ-mutant p53 interaction enhanced by Tam, thereby disrupting p73-mutant p53 complex resulting in reactivation of p73 and transcriptomic reprogramming converging on apoptosis and ferroptosis. These studies demonstrate the potential for repurposing Tam with Doxo as an effective and safe therapeutic strategy to treat molecularly stratified TNBC.

**SIGNIFICANCE:** This study demonstrates synergy between Tam and Doxo in targeting a novel ERβ-mutant p53-p73 axis in TNBC to induce apoptosis, ferroptosis, and tumor regression. These results provide a mechanistic basis for repurposing Tam in combination with Doxo as an effective therapeutic strategy against TNBC, majority of which expresses mutant p53.

## INTRODUCTION

About 15-20% of breast cancers are classified as triple negative (TNBC) (1,2), defined clinically as lacking expression of estrogen receptor alpha (ERα), progesterone receptor (PR), and human epidermal growth factor receptor 2 (HER2) overexpression. Therapeutic options to treat TNBC are limited and the current standard of care consists of chemotherapy regimens such as anthracyclines and taxanes with an increasing role for antibody-drug conjugates (ADCs), immunotherapy, and drugs targeting DNA damage and repair. TNBC metastasizes aggressively and has poor prognosis with therapeutic response rates leading to shorter survival than non-TNBCs (3,4). Overall, the current therapies have limited efficacy and could have serious adverse effects for patients. Therefore, there is an urgent need to develop new, rationally designed, therapeutic strategies.

A key molecular characteristic of TNBC is the high frequency (80%) of p53 mutation (2,5,6). Somatic mutations in p53 are very frequent and are clonally dominant, compared to other genes in TNBC (2,7). In addition to losing tumor suppressor properties and exerting dominant-negative regulation over any remaining wild type p53 (WTp53), certain mutant p53s are known to acquire oncogenic gain-of-function (GOF) properties (8–10) (11). Cellular functions affected by mutant p53 include increased invasiveness, angiogenesis, abnormal epigenetic regulation, enhancement of “stemness” and therapeutic resistance (8,11–18). p53-positivity by immunohistochemistry (an approximate surrogate for mutant p53) and p53 mutation determined by sequencing were correlated with poor survival and adverse prognostic factors in TNBC patients (19–24).

Although TNBC lacks expression of ERα, estrogen receptor beta (ERβ) is expressed in majority of these tumors (25–27). However, the role of ERβ in breast cancer has been elusive (28,29). In ERα-positive breast cancer cells, ERβ negatively regulates functions of ERα (30,31). In the case of TNBC, both anti-tumorigenic and pro-tumorigenic functions were attributed to ERβ (26) and ERβ isoforms have been shown to have contrasting activities in TNBC (32). Only some of the divergent effects could be attributed to expression of specific isoforms and their cellular location (33). Cellular signaling context and interaction with other proteins have been proposed as a plausible mechanism. Although bi-faceted functioning has been proposed to reconcile disparate findings on ERβ functions (34,35), the mechanistic basis for such duality remained largely unknown until the discovery of direct binding and functional crosstalk between ERβ and p53(36–38). Until then wild type *versus* mutant status of p53 in tumors was not factored into interpretation of ERβ function in different physiological and pathological contexts. We have shown that the capability of ERβ to function as a pro-proliferation/pro-tumorigenic protein versus anti-proliferation/anti-oncogenic protein is dependent on its interaction with wild type versus mutant p53, respectively (37). This finding was corroborated by data from our retrospective analysis of METABRIC (Molecular Taxonomy of Breast Cancer International Consortium) cohort.

As TNBC lacks expression of ERα, the canonical target of Tam, it has not been used for therapy against these cancers. Our data challenges this paradigm. We have demonstrated that Tam enhances ERβ-mediated sequestration of p73-bound mutant p53, thereby disrupting p73-mutant p53 complex resulting in reactivation of p73, leading to apoptosis and inhibition of proliferation (37). Importantly, we recently reported therapeutic proof of principle for this phenomenon in a patient with ERβ-expressing metastatic TNBC who had significant tumor regression following treatment regimen containing Tam (39).

Although Tam’s role as a SERM in cells and tumor tissues expressing ERα has been analyzed extensively, its cellular effects beyond regulating transcriptional functions of ERα-related endocrine therapy resistance has received much less attention (40–47). Here, we report for the first time that Tam synergizes with doxorubicin (Doxo), a widely used chemotherapeutic agent that inhibits topoisomerase-2, causes DNA double strand breaks, and chromatin damage by histone eviction (48). We show that Tam lowers the IC50 of Doxo and the combination therapy results in increased cell death by both apoptosis and ferroptosis in TNBC cells in vitro and suppresses tumor growth in TNBC cell line-derived xenografts (CDXs) and patient tumor-derived xenografts (PDXs). Mechanistically, along with promoting ERβ interaction with mutant p53 and reactivation of p73, Tam and Doxo combination elicits a unique pattern of transcriptome reprogramming that impinges on multiple signaling pathways that converge on cell death by ferroptosis and apoptosis. Ferroptosis is a regulated cell death process dependent on lipid peroxidation that can be exploited to kill cancer cells that are therapeutically resistant (49–52). Recent studies in cancer cell lines and mouse models have revealed important roles for mutant p53 in suppressing ferroptosis and apoptosis in cancer (9,52–54).

Our data uncover a synergy between Tam and Doxo in targeting a novel ERβ-mutant p53-p73 axis in TNBC to induce apoptosis, ferroptosis, and tumor regression. These results provide a mechanistic basis for repurposing Tam in combination with Doxo as an effective and safe therapeutic strategy against TNBC, majority of which expresses mutant p53.

## RESULTS

### 4-OHT, but not PHTPP, DPN, and 17β-estradiol, Enhances ERβ-mutant p53 Interaction in TNBC Cells

Overexpression of ERβ was reported to increase the efficacy of selective estrogen receptor modulators (SERMs) such as Tam in ERα-positive/WTp53 breast cancer cells resulting in apoptosis and cell cycle arrest (55). TNBC does not express ERα; however, majority of TNBCs express ERβ. Furthermore, ERα-positive breast cancer predominantly harbors WTp53, whereas about 80% of TNBCs express mutant p53. Following up on our previous finding that ERβ can physically interact with mutant p53 in TNBC tumors and cell lines leading to tumor suppressive effects (44,56), we embarked on understanding the mechanistic basis for these effects and exploiting them for new therapeutic strategies against TNBC. As a first step toward this goal, we analyzed the effect of ER antagonists and agonists on ERβ-mutant p53 interaction in MDA-MB- 231/LM2-4^LUC+^ (a lung metastatic variant of MDA-MB-231)(57) (hereafter termed MDA231/LM2-4 for brevity) TNBC cell line that expresses both ERβ-mutant p53. Expression of ERβ (Fig.1A), but not mutant p53 (Fig. 1B) was increased in response to treatment with 4-Hydroxy tamoxifen (4- OHT), and ERβ levels steadily increased over time in response to Tam during a time-course experiment for 0-48 hours (Fig. 1C). However, PHTPP, a synthetic non-steroidal selective ERβ antagonist at higher concentrations decreased ERβ levels (Fig. 1D), without any effect on mutant p53 levels (Supplementary Fig. S1A). DPN, a synthetic non-steroidal ERβ-specific agonist as well as 17β-estradiol (E2), a potent agonist for ERα and ERβ, did not affect expression levels of ERβ (Fig. 1E and Supplementary Fig. S1F) and mutant p53 (Supplementary Fig. S1B). Proximity ligation assay (PLA) for analyzing *in situ* interaction between ERβ and mutant p53 showed that the interaction was increased by 4-OHT (Fig.1F), whereas PHTPP decreased the interaction (Fig. 1G). Neither DPN (Fig. 1H) nor 17β-estradiol (E2) (Supplementary Fig. S1D) affected the interaction. Single antibody was used as negative control (Supplementary Fig. S1E). Consistent with the PLA data, co-immunoprecipitation assay revealed higher levels of ERβ-mutant p53 interaction in cells treated with 4-OHT (Fig.1I). Cell treated with PHTPP had lower levels of interaction (Fig. 1 J) while DPN treatment did not considerably affect the interaction (Fig. 1K). These data demonstrate the ability of Tam to increase ERβ protein levels and increase ERβ- mutant p53 interaction in TNBC cells.

**Figure 1.**
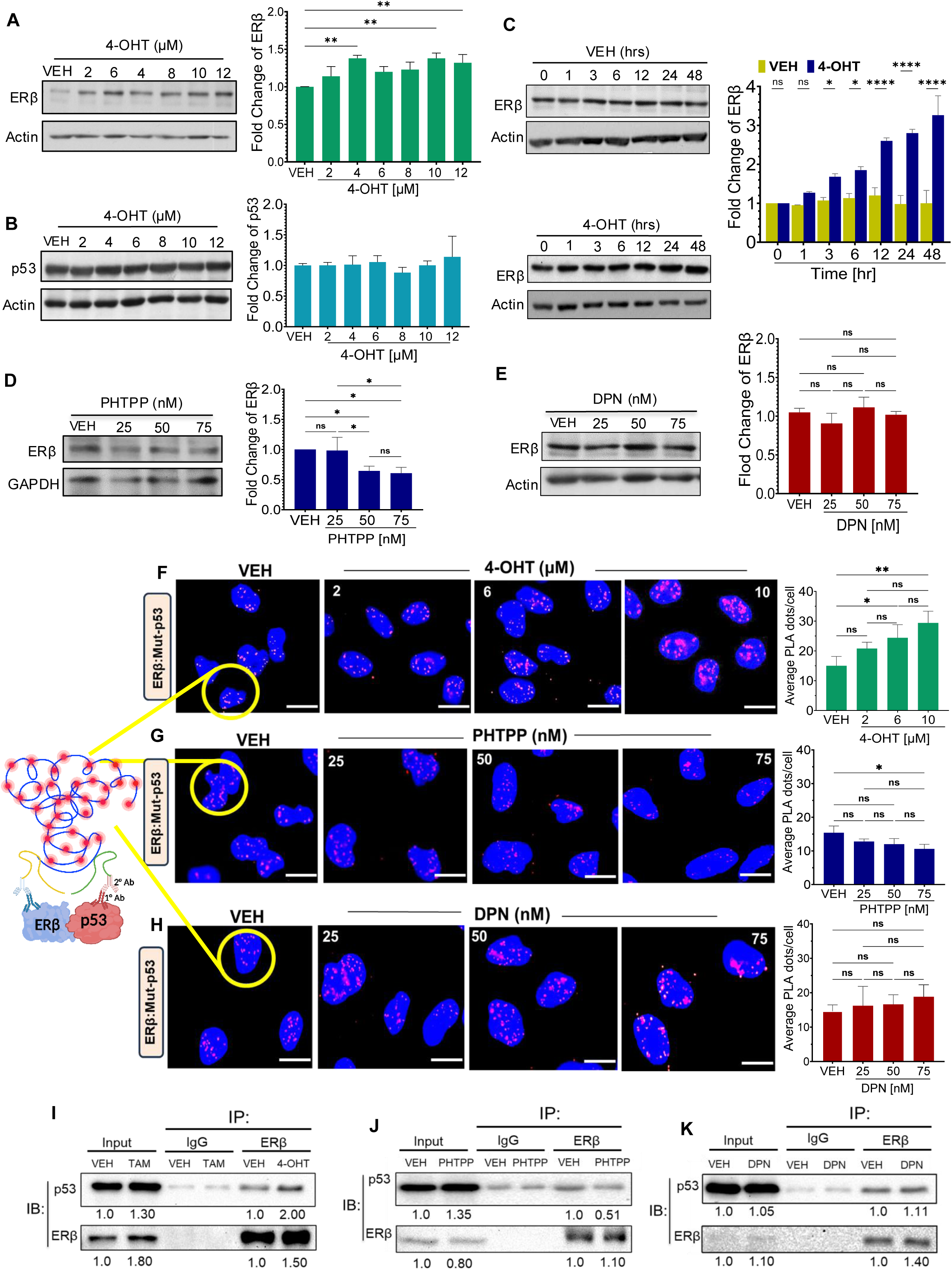
4-Hydroxytamoxifen stabilizes ERβ and enhances its interaction with mutant p53 in TNBC cells. A and B, MDA-231/LM2-4 cells were treated with vehicle (Veh) or increasing concentrations of 4- hydroxytamoxifen (4-OHT; 2,6,4,8,10 &12 μM) for 24 h. Post treatment, cells were lysed in radioimmunoprecipitation assay (RIPA) buffer Immunoblot analysis of ERβ **(A)** and p53 **(B)** protein levels is shown, with β-actin serving as the loading control. Right, densitometric quantification of ERβ and p53 expression normalized to β-actin. **C,** Cells were treated with Veh (top panel) or 4- OHT (10 μM (bottom panel) for 0, 1, 3, 6, 12, 24, or 48 h. ERβ protein expression was assessed by immunoblotting at the indicated time points (left) and densitometric quantification of ERβ protein levels normalized to β-actin, which served as the loading control (right). **D & E,** Immunoblot analysis of ERβ protein expression following 24 h of treatment with increasing concentrations of the ERβ antagonist PHTPP (25, 50, and 75 nM) **(D)** or the ERβ agonist DPN (25, 50, and 75 nM) **(E).** Densitometric quantification of ERβ protein levels normalized to β-actin (right panels). **F–H,** MDA-MB-231/LM2-4 cells were treated for 24 h with Veh or increasing concentrations of 4-OHT (2, 6, & 10 μM) **(F)**, ERβ antagonist PHTPP (25, 50, and 75 nM) **(G)**, or the ERβ agonist DPN (25, 50, and 75 nM) **(H)**. Following treatment, cells were fixed with 4% paraformaldehyde and subjected to proximity ligation assay (PLA) to evaluate the physical interaction between ERβ and mutant p53 proteins. Representative PLA images are shown in the left panels, where red fluorescent puncta indicate ERβ-mutant p53 interaction events. Nuclei were counterstained with DAPI (blue). The right panels show quantitative analysis of PLA signals, expressed as the average number of PLA puncta (dots) per cell. Quantification was performed using ten selected microscopic fields per condition from three independent biological experiments. **I–K,** Co- immunoprecipitation (co-IP) analysis of the interaction between ERβ and mutant p53 following 24 h treatment with Veh, 4-OHT (2 μM) **(I),** PHTPP (50 nM) **(J),** or DPN (50 nM) **(K),** compared with untreated controls. Fold changes in ERβ and p53 expression in ligand-treated samples were normalized to their respective vehicle-treated controls. Densitometric analysis of ERβ and p53 was performed using ImageJ, and all quantitative data were plotted and analyzed in GraphPad Prism 9 (**A-E**). Data in **(A–H)** are presented as mean ± SD from three independent experiments. Statistical significance was assessed using one-way ANOVA for Fig. **A, B, & D–H**, and Student’s *t*-test for Fig. **C**. Immunoblots in Fig. **I–K** show representative results from single experiment for each drug treatment. Statistical significance is indicated as follows: ns, not significant; *P ≤ 0.05; **P ≤ 0.01; and ****P ≤ 0.0001.

### Mutant p53 is Necessary for Tamoxifen-Mediated Anti-Proliferation and Anti-Tumorigenic Effects in TNBC Cells

Consistent with its ability to increase ERβ-mutant p53 interaction, 4-OHT upregulated mRNA (Fig. 2A) and protein (Fig. 2C) levels of p21 *(CDKN1A*) and PUMA (*BBC3*) in MDA231/LM2 cells with endogenous mutant p53 (p53R280K), whereas PHTTP downregulated the expression of these genes (Fig. 2A and 2C). However, such increase in RNA or protein levels were absent in the isogenic MDA231/LM2-4^LUC+^p53KO cells where mutant p53 was knocked out (Fig. 2B and 2D). DPN did not affect expression of *CDKN1A* and *BBC3* in MDA231/LM2-4^LUC+^ (Supplementary Fig. 2A) or MDA231/LM2-4^LUC+^p53KO cells (Supplementary Fig. 2B). Consistent with these data, 4- OHT increased G1 phase arrest (BrdU assay) (Fig. 2E and Supplementary Figure S2C) and apoptosis (annexin V-positive staining) (Fig. 2F and Supplementary Figure S2E) in cells expressing mutant p53, but not in p53KO cells (Fig. 2G, 2H, and Supplementary Figure S2D and S2F). Similar results were obtained in another TNBC cell line MDA-MB-468 expressing a different mutant p53 (p53R273H) (Fig. 2I, 2J, 2K, Supplementary Figure S2G and S2H). treatment with PHTTP decreased G1 arrest in MDA231/LM2-4^LUC+^ cells (Supplementary Fig. 2I), but not in MDA231/LM2-4^LUC+^KO cells (Supplementary Fig. 2J). DPN did not have any effect on cell clycle in MDA231/LM2-4^LUC+^ (Supplementary Fig. 2K), MDA231/LM2-4^LUC+^KO (Supplementary Fig. 2L), and MDA-MB-368 (Supplementary Fig. 2L) cells. RNA-seq analysis of gene expression in MDA231/LM2-4^LUC^ cells treated with Tam revealed that genes encoding *BTG2*, a p53 target that is involved in cell differentiation, DNA damage repair, apoptosis, and negatively regulating TNBC tumor growth (58,59); *PUMA/BBC3*, a major pro-apoptotic protein; *E2F7*, an atypical E2F family member that restricts homologous recombination and has a repressive role in DNA damage response (DDR) (60); and *TIMP2,* a suppressor of TNBC growth and metastasis through modulation of EMT, vasculature, and metastasis signaling pathways (61) were upregulated (Fig. 2L). However, when mutant p53 was knocked out (p53KO cells), these genes were no longer activated in response to Tam (Fig. 2M). Instead, Tam downregulated *BTG2, BBC3*, and *E2F7* in p53KO cells. Furthermore, genes encoding pro-metastasis proteins such as CXCL1 (62) and LSM4 (63) were down regulated in MDA231/LM2-4^LUC+^ cells (Fig. 2L). On the contrary, Tam upregulated *CXCL1* in MDA231/LM2-4^LUC+^ p53KO cells (Fig. 2M). These data suggest that antiproliferation effect of Tam in TNBC is mostly dependent on mutant p53 expression.

**Figure 2.**
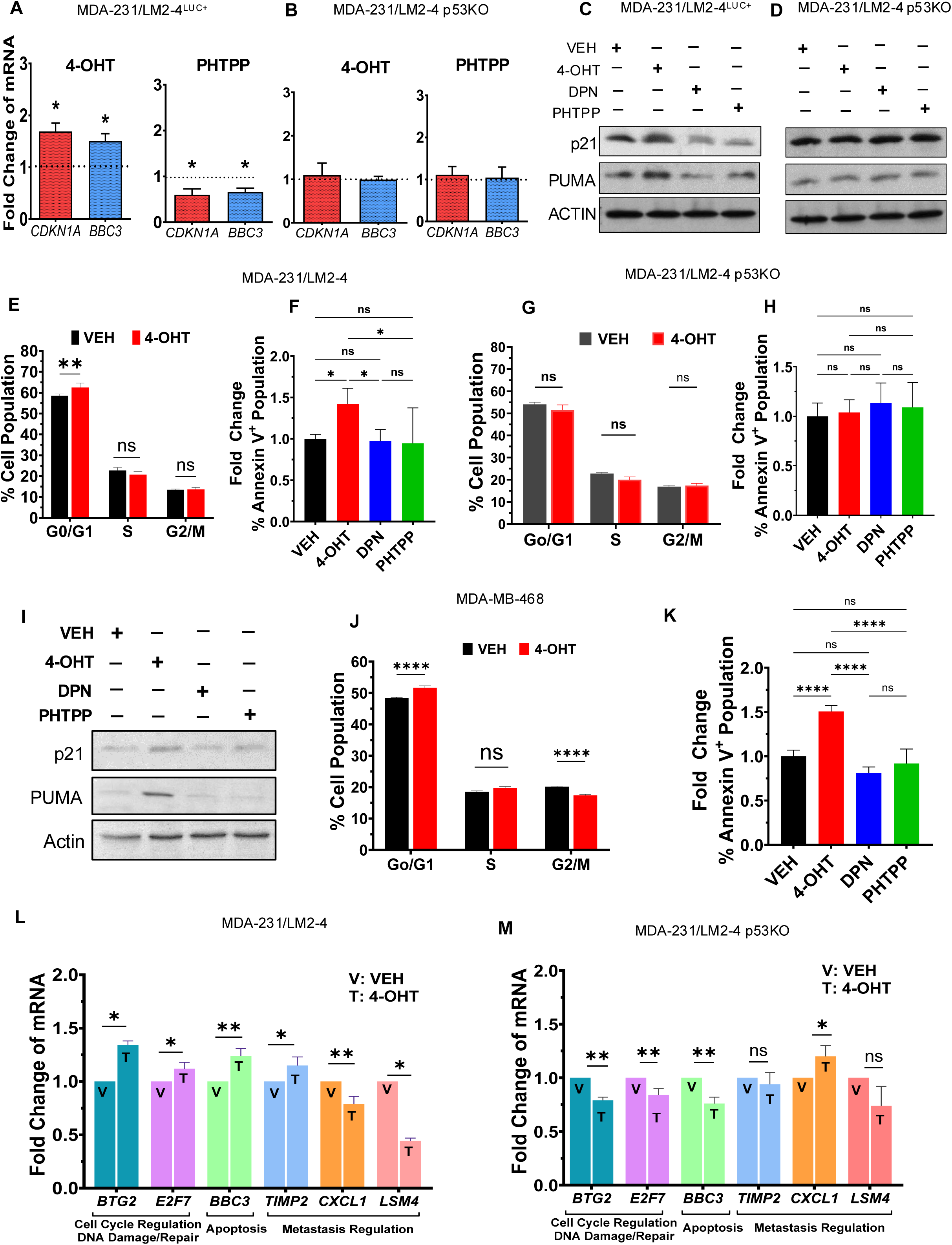
Mutant p53 is necessary for anti-tumorigenic effects of tamoxifen in TNBC cells A and. **B**, MDA-231/LM2-4 (**A**) and isogenic MDA-231/LM2-4 p53-knockout (p53KO) (**B**) cells were treated with Veh, 4-OHT (2 μM), or PHTPP (50 nM) for 24 hours. Post treatment, total RNA was isolated, and transcript levels of *CDKN1A* (p21) and *BBC3* (PUMA) were quantified by quantitative real-time polymerase chain reaction (qRT–PCR). **C–D,** MDA-231/LM2-4^LUC+^ (C) and p53KO (D) cells were treated with VEH, 4-OHT (2 μM), DPN (50 nM), or PHTPP (50 nM) for 24 h. Whole-cell lysates were subjected to immunoblot analysis for p21 and PUMA, with β-actin as the loading control. **E,** MDA-MB-231/LM2-4 cells were treated with Veh, 4-OHT; 10 µM for 24 h, followed by a 1-h pulse with 10 µM BrdU. Cell-cycle distribution was assessed by flow cytometry. **F,** MDA-MB- 231/LM2-4 cells were treated with Veh and 4-OHT (10 µM) for 24 h. Apoptosis was assessed by Annexin V–FITC/propidium iodide (PI) staining. Percent positive cells were analyzed by flow cytometry. **G & H,** p53KO cells were treated with VEH or 4-OHT (10 µM) for 24 h. Post treatment cells were pulsed with 10 µM BrdU for 1 h for cell-cycle distribution analysis **(G)** and stained with annexin V–FITC/PI, followed by flow cytometry for apoptosis analysis **(H)**. **I,** MDA-MB-468 cells were treated with VEH, 4-OHT (2 μM), DPN (50 nM), or PHTPP (50 nM) for 24 hours. Post treatment, cells were lysed in RIPA buffer and immunoblotted for p21 and PUMA with β-actin as a loading control. **J & K,** MDA-MB-468 cells were treated with VEH or 4-OHT (10 µM) for 24 h. Cells were pulsed with 10 µM BrdU for 1 h for cell-cycle distribution analysis **(J)** and stained with annexin V–FITC/PI for apoptosis assessment **(K),** followed by flow cytometric analysis. **L and M,** MDA-MB-231/LM2-4 **(L)** and p53KO cells **(M)** were treated with Veh or 4-OHT; 2 μM for 24 h, followed by next-generation RNA sequencing (RNA-seq). Relative transcript levels of *BTG2, E2F7, BBC3, CXCL1, TIMP2,* and *LSM4* are shown based on differential gene expression analysis normalized with Veh-treated group. Data shown in Fig. A, B, E**–H**, and **J–M** represent three independent experiments, whereas Fig. **C** and **D** represent a single experiment and Fig. **I** represents two independent experiments. Quantitative data were analyzed and plotted using GraphPad Prism 9. Data is presented as mean ± SD. Statistical significance between two groups was determined using Student’s *t*-test for Fig. A, B, E, G**, J, L,** & **M**. For comparisons among multiple treatment groups, one-way ANOVA was used for Fig. F, H, and **K**. Statistical significance is indicated as follows: ns, not significant; *P ≤ 0.05; **P ≤ 0.01; and ****P ≤ 0.0001.

### 4-OHT Synergizes with Doxorubicin to Induce Apoptosis and to Suppress Tumor Cell Migration and Invasion of TNBC Cells

We used Chou-Talalay method (64) to analyze whether 4-OHT combined with Doxo produces additive or synergistic effect in MDA231/LM2-4^LUC+^ cells. The analysis performed with 4-OHT and different concentrations of Doxo showed that addition of 4-OHT lowers the IC50 of Doxo in the MDA231/LM2-4^LUC+^ cells with endogenous R280K mutant p53 expression (Fig. 3A, Supplementary Fig. S3B and S3C). The combination index (CI) was <1 suggesting the effect was synergistic (Fig. 3B). However, synergistic effect was not observed in the isogenic cell line MDA231/LM2-4^LUC+^ p53KO where mutant p53 was knocked out (KO) (Fig. 3C), suggesting that mutant p53 is necessary for synergy between Tam and Doxo. Furthermore, synergy was observed in in MDA-MB-468 TNBC cells expressing endogenous p53 R273H mutation (Supplementary Fig. S3A). Consistent with this observation, RNA-seq analysis of gene expression in MDA231/LM2- 4^LUC+^ and MDA231/LM2-4^LUC+^ p53KO cells treated with Tam, Doxo, or Tam+Doxo combination showed that expression of pro-apoptotic genes including *TNFRSFB10B* (encodes DR5), *CASP8* (encodes caspase 8), both transcriptional targets of WTp53, were highly upregulated in response to combination therapy compared to monotherapy with either drug in the cells expressing mutant p53, but not in p53KO cells (Fig. 2D). The expression of *TNFRSF10B*, and *CASP8* were ascertained by qRT-PCR (Fig. E and Fig. F). Furthermore, the levels of apoptosis protein markers, cleaved PARP and Caspase, were upregulated in response to Tam+ Doxo combination as compared to individual drugs in MDA231/LM2-4^LUC+^ cells (Fig. 3G). In the case of p53KO cells, cleaved PARP and caspase protein levels were increased in response to both Doxo treatment and Doxo+Tam combination (Fig. 3H). This response is apparently due to active p73 in the absence of mutant p53 in these cells. These data were corroborated by the finding that there was increased apoptosis (Annexin V assay) in response to Tam + Doxo treatment (as compared to treatment with individual drugs) of cells expressing mutant p53 (Fig. 3I) and is consistent with changes in cellular morphology (Supplementary Fig. S3E and S3F). Again, the increased ability of the drug combination to induce apoptosis was compromised in p53KO cells (Fig. 3J).

**Figure 3.**
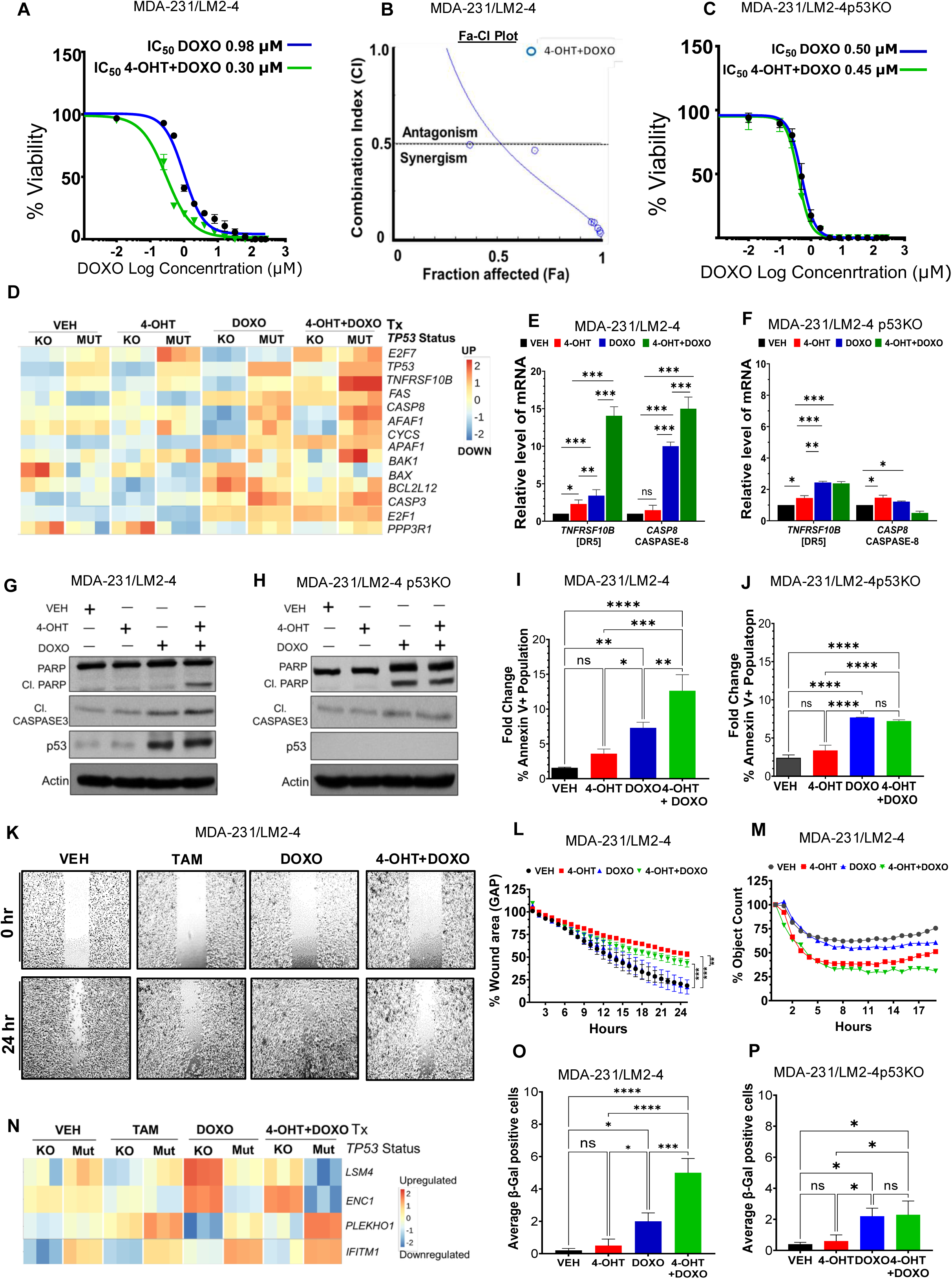
4-Hydroxytamoxifen synergizes with doxorubicin to induce apoptosis and suppresses cell proliferation, migration, and invasion in TNBC cells. **A**, MDA-MB-231/LM2-4 cells were treated with increasing concentrations of DOXO (0.001, 0.01,0.1, 0.25, 0.5, 1, 2, 4, 8, &12 μM), either alone or in combination with 4-OHT (10 μM), for 48 h . Cell growth inhibition was assessed to generate dose response curves and determine the IC₅₀ values for DOXO alone and the combination treatment. **B**, The dose effect relationship of 4-OHT and DOXO, administered individually or in combination at a fixed 1:1 ratio, was analyzed using CompuSyn software. Combination index (CI) plot showing the CI values across different fractional effects. CI values were calculated using the Chou–Talalay method, where CI < 1 indicates synergism, CI = 1 indicates an additive effect, and CI > 1 indicates antagonism. **C,** p53KO cells were treated with increasing concentrations of DOXO, either alone or in combination with 4-OHT, as described in **A**. Cell growth inhibition was assessed to generate dose response curves and determine the IC₅₀ values for DOXO alone and the combination treatment. **D,** Heatmap showing differentially expressed apoptosis-related genes identified by RNA-seq in parental MDA-MB- 231/LM2-4^LUC⁺^ (MUT) and p53KO (KO) cells following treatment with Veh, 4-OHT; 10 μM, DOXO; 0.5 μM, or the combination of 4-OHT and DOXO (0.5 μM) for 24 h. **E&F**, qRT-PCR analysis of *TNFRSF10B* (DR5) and *CASP8* (caspase-8) gene expression in MDA-MB-231/LM2-4 **(E)** and p53KO **(F)** cells following treatment with Veh, 4-OHT; 10 μM, DOXO; 0.5 μM, or the combination for 24 h. **G&H,** Immunoblot analysis of full-length PARP, cleaved PARP, cleaved caspase, p53, and β-actin in MDA-MB-231/LM2-4 **(G)** and p53KO **(H)** cells following treatment with Veh,4-OHT; 10 μM, DOXO; 0.5 μM, or the combination for 24 h. β-actin was used as a loading control. **I &J**, MDA-MB-231/LM2-4 **(I)** and p53KO **(J)** cells following treatment with Veh, 4-OHT; 10 μM, DOXO; 0.1 μM, or the combination for 24 h. Apoptosis was assessed by Annexin V– FITC/propidium iodide (PI) staining. Percent positive cells were analyzed by flow cytometry. **K**, Representative phase-contrast images of wound-healing assays showing cell migration in MDA- MB-231/LM2-4 cells treated with Veh, 4-OHT; 10 μM, DOXO; 0.5 μM, or the combination of 4- OHT (10 μM) and DOXO (0.5 μM). Images were captured at 0 and 24 h to assess wound closure over time. Scale bar, 100 μm. **L**, Quantitative analysis of wound closure in MDA-MB-231/LM2-4 cells treated as described in (**K)**. Wound closure was quantified from 0 to 24 h to assess the effects of individual and combination treatments on cell migration. **M**, Quantitative analysis of Matrigel invasion of MDA-MB-231/LM2-4 cells following treatment with Veh, 4-OHT; 10 μM, DOXO; 0.1 μM, or the combination for 24 h. Invasion was assessed over 0–20 h to evaluate the effects of individual and combination treatments on the invasive capacity of the cells. **N**, Heatmap of differentially expressed genes identified by RNA-seq highlighting selected pro-metastatic (*LSM4 & ENC1*) and anti-metastatic genes (*PLEKHO1 & IFITM1*) in MDA-MB-231/LM2-4 (Mut) and p53KO (KO) cells following treatment with Veh, 4-OHT; 10 μM, DOXO; 0.5 μM, or the combination for 24 h. The heatmap illustrates relative gene expression patterns across treatment groups. **O&P,** MDA-MB-231/LM2-4 **(O)** and p53KO **(P)** cells treated with 4-OHT; 10 μM, DOXO; 0.5 μM, or the combination for 24 h. Following treatment, cells were incubated with the β- galactosidase staining reagent for 12 h. SA-β-gal-positive cells were identified and quantified using bright-field microscopy. Representative bright-field microscopy images of SA-β-gal staining are shown in Supplementary Fig. **S3I** and **S3J**. Quantitative data in panels **A-l & N-P** are as mean ± SD from three independent experiments. Statistical significance was assessed using one-way ANOVA. Statistical significance is indicated as follows: ns, not significant; \**P* ≤ 0.05; **P ≤ 0.01; \*\*\**P* ≤ 0.001; and ****P ≤ 0.0001. Data in panel **M** are from a single experiment.

Importantly, Tam as well as Tam + Doxo treatment reduced both migration (Fig. 3K and 3L) and invasion (Fig. 3M) of MDA231/LM2-4^LUC+^ cells, but not of the isogenic p53KO cells (Supplementary Fig. S3G and S3H). Consistent with these data, RNA-seq analysis showed that expression of pro-metastatic genes such as *LSM4,* an oncogenic RNA-binding protein promoting TNBC cell proliferation, migration, and invasion (65), and *ENC1,* an inducer of metastasis in breast cancer by promoting Wnt/β-catenin pathway (66), are downregulated by Tam + Doxo combination (Fig. 3N). On the other hand, metastasis suppressor genes *PLEKHO1*(*CKIP-1*) (67) and *IFITM1* ((68) are upregulated in response to Tam+Doxo combination in mutant p53- expressing MDA231/LM2-4^LUC+^ cells, but not in the isogenic p53KO cells (Fig. 3N).

In addition to Tam+Doxo combination effectively in upregulating apoptosis, and downregulating cell migration and invasion in TNBC, it increased cellular senescence as well in cells with mutant p53 (Fig. 3O) and (Supplementary Fig. S3I), but not as much in MDA231/LM2-4^LUC+^p53KO cells (Supplementary Fig. S3J); however, the combination treatment had no advantage over Doxo alone treatment in upregulating senescence (Fig. 3P) and (Supplementary Fig. S3J). Together, these data demonstrate that Tam+Doxo combination strong antitumor effects against TNBC expressing mutant p53 by activating apoptosis and inhibiting cell migration and invasion.

### Estrogen Receptor β Sequesters p53 with Hot-Spot Mutations Leading to Enhanced Apoptosis in Response to Combination Therapy

p53 is mutated in about 80% of TNBC and different missense mutations, especially in the DNA-binding domain, have been reported. For analyzing the effect of different p53 contexts on therapeutic response in TNBC cells, we generated isogenic MDA231/LM2-4^LUC+^ cell lines with (i) endogenous R280K mutant p53 knocked out (p53 KO) without or with ERβ knockdown, (ii) stable expression in p53KO cells, of WT p53 and different p53 mutants (R175H, R273H, R248Q and R280K) without or with ERβ knockdown (Supplementary Fig. S4A). We analyzed the effect of these different p53 contexts on ERβ-p53 interaction and on eliciting apoptosis in response to 4- OHT and Doxo monotherapies *versus* 4-OHT + Doxo combination therapy. PLA demonstrated interaction of ERβ with p53 with R175H, R273H, R248Q hot-spot mutations and the endogenous R280K mutation that is expressed in parental MDA231/LM2-4^LUC+^ cells, whereas Interaction with wild type p53 was relatively low (Fig. 4A and 4B). R175H and R280 K mutants showed higher binding affinity (lower *K_D_*) to ERβ than WTp53 (Supplementary Fig. 4B). Isogenic cell lines expressing mutant p53 showed lower IC_50_ for 4-OHT + Doxo combination as compared to monotherapy with Doxo (Fig. 4C, bar pairs 2-5, and Supplementary Fig. 4C), whereas cells expressing wild type p53 had the opposite response, i.e., higher IC_5o_ for combination therapy (Fig. 4C, bar pair 6). Knocking down endogenous ERβ in MDA231/LM2-4^LUC+^ cells (expressing mutant p53R280K) resulted in higher IC_50_ for the combination therapy reversing the trend in cells expressing mutant p53 and ERβ, demonstrating that ERβ is necessary for synergistic activation effect of 4-OHT + Doxo combination. Similarly, knocking out mutant p53 in MDA231/LM2-4^LUC+^ cells expressing endogenous ERβ (MDA231/LM2-4^LUC+^p53KO cells) did not show lower IC50 for the combination treatment as compared to monotherapy with Doxo. These data strongly suggest that mutant p53 and ERβ physical interaction is required for increased responsiveness (decreased IC_50_) to 4-OHT + Doxo combination.

**Figure 4.**
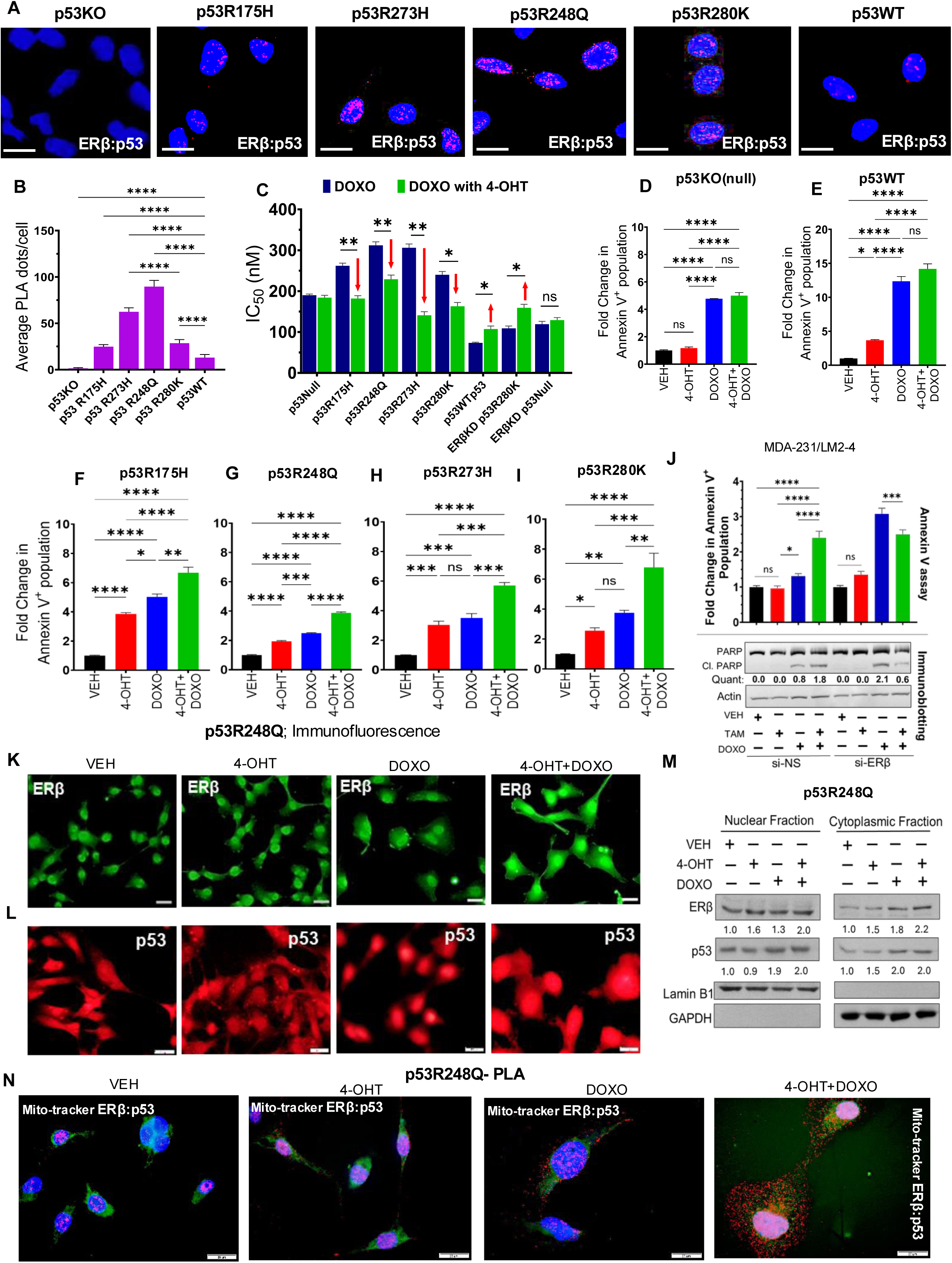
ERβ interaction with gain-of-function p53 mutants enhances apoptotic response to 4- Hydroxy tamoxifen and doxorubicin combination treatment. **A**, Isogenic MDA-MB-231/LM2-4 cells p53KO, wild-type p53 (p53WT), or the mutant p53 variants R175H, R248Q, R273H, and R280K were analyzed by PLA to assess basal interactions between ERβ and p53. Representative fluorescence microscopy images showing PLA signals (pink dots), indicative of ERβ–p53 interaction are shown. Scale bar, 10 μm. **B,** Quantification of PLA interaction signals were from 10 independent microscopic fields per condition across three independent experiments. **C,** Isogenic cells expressing p53KO, p53WT, & p53 variants (R175H, R248Q, R273H, and R280K), as well as ERβ-knockdown cells expressing p53R280K (ERβKD/p53R280K) and ERβ- knockout cells with p53KO (ERβKD/p53KO) were treated with increasing concentrations of DOXO (0.001, 0.01, 0.1, 0.25, 0.5, 1, 2, 4, and 8 μM), either alone or in combination with 4-OHT; 10 μM, for 48 h. Cell growth inhibition was measured, and dose response curves were generated to determine the IC₅₀ of DOXO alone and in combination with 4-OHT. **D-I,** Quantification of apoptosis in the isogenic TNBC cell models p53KO **(D),** p53WT **(E)**, p53R175H **(F)**, p53R248Q **(G)**, p53R273H **(H)**, and p53R280K **(I)** following treatment with Veh, 4-OHT; 10μM, DOXO; 0.1 μM, or the combination for 24 h. Following treatment, cells were stained with Annexin V and the percentage of Annexin V- positive cells was quantified to assess treatment-induced apoptosis. Quantitative data were plotted using GraphPad Prism 9. **J,** MDA-MB-231/LM2-4 cells were transiently transfected with non-specific control siRNA (si-NS) or ERβ specific siRNA (siERβ) for 48 h, followed by treatment with Veh, 4- OHT; 10 μM, DOXO; 0.1 μM, or the combination for 24h. Post treatment cells were stained with Annexin V^+^ and positive cells were analyzed by flowcytometry (Top). For immunoblot analysis, post treatment cells were lysed by RIPA buffer and PARP and cleaved PARP levels were analysed, with β-actin serving as the loading control (bottom). Cleaved PARP levels were quantified and normalized to β-actin. **K & L**, Representative immunofluorescence microscopic images showing expression of ERβ (green, **K**) and mutant p53 (red, **L**) in the isogenic cell line expressing p53R248Q following treatment with 4-OHT (10 μM), DOXO (0.5 μM), or the combination for 24 h. Scale bar;10μm. **M,** Immunoblot analysis of nuclear and cytoplasmic fractions of p53R248Q isogenic cells following the treatment of Veh, 4-OHT (10 μM), DOXO (0.5 μM), or the combination for 24 h. Quantification of nuclear and cytoplasmic expression of ERβ and p53 is normalized to nuclear fraction loading control LaminB1 and cytoplasmic loading control GAPDH respectively. **N,** Isogenic p53R248Q cells were treated with VEH, 4-OHT (10 µM), DOXO (0.5 µM), or the combination for 24 h, Post treatment cells were fixed with live Mito tracker (Green) stain for 1 hour in CO2 incubator at 37⁰C. Post incubation cells were fixed with 2% formaldehyde followed by PLA analysis of ERβ and p53 interaction. Representative PLA (Pink), DAPI (blue) with merged Mito-tracker (Green) are shown. Scale bar;10 µm. Data in Fig. **B-L&N** are presented as mean ± SD from three independent experiments. Statistical significance was assessed using one-way analysis of variance (ANOVA) for Fig. **B and D–J**, and Student’s *t*-test for Fig. **C**. Statistical significance is indicated as follows: ns, not significant; \**P* ≤ 0.05; \*\***P** ≤ 0.01; \*\*\**P* ≤ 0.001; and \*\*\*\***P** ≤ 0.0001. Data in Fig. **M**, is representation of two independent experiments.

Consistent with these IC50 data, 4-OHT + Doxo combination did not have any advantage over monotherapy with Doxo in inducing apoptosis (Annexin V assay) of MDA231/LM2-4^LUC+^p53KO cells expressing endogenous ERβ and lacking mutant p53 (Fig. 4D, and Supplementary Fig. S4D). On the contrary, apoptosis in response to 4-OHT + Doxo combination was much stronger as compared to response to either Tam or Doxo monotherapies in all the isogenic cell lines expressing mutant p53 (Fig. 4E-I). However, advantage of 4-OHT + Doxo combination over Doxo monotherapy in inducing apoptosis (assayed by Annexin V staining and PARP cleavage) in MDA231/LM2-4^LUC+^ cells expressing endogenous mutant p53R280K, while this advantage was lost when ERβ was knocked down (Fig. 4J). These data provide further support for the requirement of both mutant p53 and ERβ for the cells to be responsive to 4-OHT + Doxo combination.

Importantly, levels of ERβ mutant p53 were increased in nuclear and cytoplasmic compartments in response to treatment (Fig. 4K, 4L, 4M, and Supplementary Fig. 4E). Of note, PLA with ERβ and p53 antibodies along with immunofluorescence with mitochondrial marker mitotracker revealed robust ERβ-mutant p53 interaction in the nucleus and cytoplasm, including mitochondria (Fig. 4N, Supplementary Fig. 4F).

These data demonstrate that TNBC cells expressing ERβ and p53 with different mutations are responsive to 4-OHT + Doxo combination therapy, and that both ERβ and mutant p53 are necessary for the increased response to 4-OHT + Doxo combination treatment over monotherapy with these agents.

### ERβ-Mutant p53 Interaction Enhances Pro-Apoptosis Function of p73 in Response to Combination Therapy

Binding and functional inactivation of tumor suppressor p73 is one of the prominent gain-of-oncogenic functions of mutant p53 (69) . We have previously reported that sequestration of mutant p53 by ERβ leads to disruption of mutant p53-p73 interaction and reactivation of p73(37). We analyzed if this phenomenon is relevant to the cellular response to 4-OHT + Doxo combination treatment in TNBC cells. As expected, PLA showed that ERβ-mutant p53 interaction is increased in response to 4-OHT monotherapy as well as 4-OHT + Doxo combination therapy (Fig. 5A, upper panel). There is corresponding decrease in p73-mutant p53 interaction in these cells except in those treated with Doxo alone (Fig. 5A, bottom panel). This increase in p73-mutant p53 interaction is likely because of increased levels of both p73 and mutant p53 proteins in response to Doxo treatment (Fig. 5I, lane 3). As ERβ level did not increase in response to Dox (Fig. 5I, lane 3), the available ERβ may not be sufficient to sequester the large amount of p53 away from p73, resulting in high p73-mutant p53 interaction. Importantly, when Doxo is combined with Tam, p73-mutant p53 interaction is decreased (Fig 5A, bottom panel, and PLA quantitation graph on the right) likely because of the high levels of ERβ sequestering away mutant p53 and keeping it unavailable to complex with p73, leading to reactivation of p73 and upregulation of cell death.

**Figure 5.**
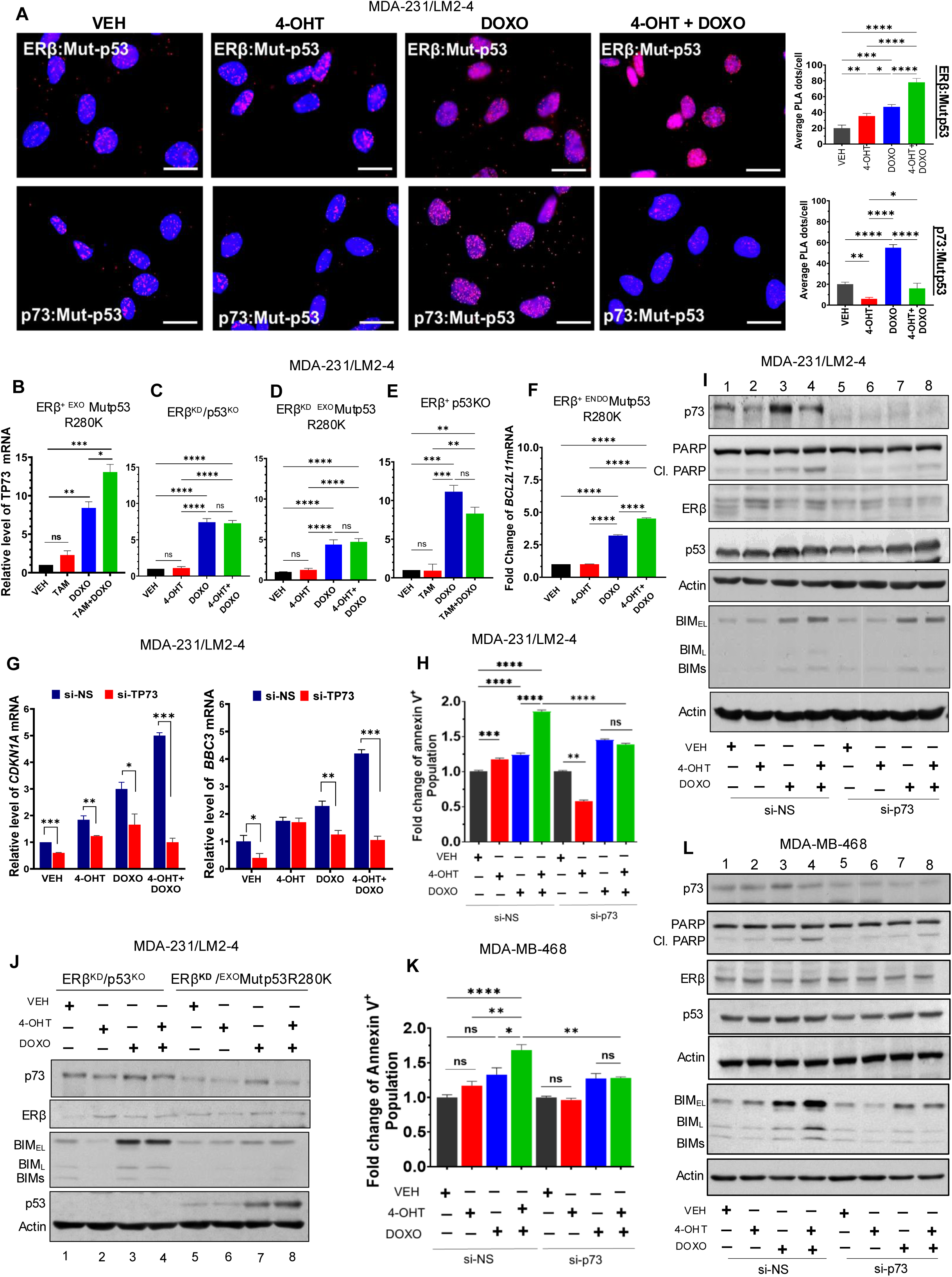
ERβ sequesters mutant p53 from p73 to enhance apoptotic response to 4-Hyrdoxy tamoxifen and doxorubicin combination treatment. A, MDA-MB-231/LM2-4 cells were treated with Veh, 4-OHT; 10 μM, DOXO; 0.5 μM, or the combination 24 h. Following treatment, cells were fixed and subjected to PLA to evaluate the physical interaction of ERβ-mutant p53 and p73-mutant p53. Representative PLA images are shown where pink, fluorescent puncta as shown. Scale bar;10 μm. Quantification of PLA signals is presented in the right panel and expressed as the average number of interaction puncta per cell. **B-F,** qRT-PCR analysis was performed to determine the differential expression of *TP73* (p73) and *BCL2L11* (BIM) following treatment with Veh, 4-OHT;10 μM, DOXO; 0.5 μM, or the combination of 4-OHT and DOXO for 24 h. *TP73* mRNA expression was evaluated in ERβ ^EXO^MUTp53R280K cells **(B),** ERβKD/p53KO cells **(C),** ERβKD^EXO^MUTp53R280K cells **(D),** and ERβ/p53KO cells (E). *BCL2L11* (BIM) mRNA expression was analyzed in MDA-231/LM2- 4^LUC^⁺^ENDO^MUTp53R280K cells **(F)**. Relative gene expression levels were normalized to the indicated housekeeping gene and expressed as fold change relative to the vehicle-treated control group. **G-I,** MDA-231/LM2-4 cells were transfected with non-specific (si-NS) or p73-targeting siRNA (si-TP73) for 48 h, followed by treatment with VEH, 4-OHT (10 µM), DOXO (0.5 µM), or combination for 24 h. Transcript levels of *CDKN1A* (p21) and *BBC3* (PUMA) were quantified by qRT-PCR **(G)**. After treatment, cells were stained with Annexin V to detect apoptotic cell death and analyzed by flow cytometry **(H)**. The percentage of Annexin V-positive cells, representing early and late apoptotic populations, was quantified and is presented in the graph. Immunoblotting expression analysis of p73, PARP, cleaved PARP, ERβ, p53, and BIM **(I). J,** Immunoblot analysis of p73, ERβ, p53, and BIM of TNBC isogenic cells expressing ERβKD/p53KO and ERβKD/p53^EXO^MUTp53R280K following treatment Veh, 4-OHT (10 μM), DOXO (0.5 μM), or the combination for 48 h. **K&L,** MDA-MB-468 cells were transfected with non-specific (si-NS) or p73- targeting siRNA (si-TP73) for 48 h, followed by treatment with VEH, 4-OHT (10 µM), DOXO (0.5 µM), or combination for 24 h. Post treatment, cells were stained with Annexin V stain and percentage cells were analyzed using flowcytometry **(K).** Immunoblot analysis of p73, PARP, cleaved PARP, ERβ, p53, and BIM **(L).** Quantitative data presented in Fig. **A–F, H,** and **K** represent the mean ± SD from three independent biological experiments and statistical analysis was performed using one-way ANOVA and Fig. **G** were analyzed using a two-tailed Student’s *t*- test. Statistical significance is indicated as follows: ns: non-significant, *P ≤ 0.05, **P ≤ 0.01, ***P ≤ 0.001, ****P ≤ 0.0001 significant. Data shown in Fig. I, J, and **L** are representative of two experiments.

In TNBC cells expressing mutant p53, the combination therapy is more efficient than monotherapies in upregulating p73 mRNA levels (Fig. 5B). In cells where both ERβ and mutant p53 are depleted, combination therapy has no advantage over Doxo monotherapy in upregulating p73 mRNA, likely because in the absence of mutant p53, p73 remains functionally active to autoregulate its own transcriptional upregulation in response to genomic damage by Doxo (Fig. 5C) (70). Furthermore, when ERβ is depleted in TNBC cells expressing mutant p53, p73 mRNA levels remain very low in response to both Doxo monotreatment as well as Tam+Doxo combination (Fig. 5D), apparently because of the absence of ERβ to bind and disable mutant p53 from suppressing functional p73, thereby resulting in lack of feed-forward upregulation of p73 transcription by p73 protein (70).

Thus, both ERβ and mutant p53 are necessary for TNBC cells to be responsive to the combination therapy. Suppression of transcriptionally active p73 by mutant p53 resulting in inhibition of autoregulated increase in p73 RNA levels is further supported by the finding that knocking out mutant p53 (p53KO cells) leads to high levels of p73 mRNA in response to both Doxo and Tam+Doxo treatments (Fig. 5E).

We addressed the functional significance of differential increase in p73 levels in MDA/231/LM2- 4Luc cells in response to Doxo versus Tam + Doxo treatments. *BCL2L11* gene encodes BIM, a major transcriptional target of p73 (71) (69,70) and pro-apoptotic protein. Tam + Doxo is more effective than either Tam or Doxo monotherapies in increasing the expression of *BCL2L11* RNA (Fig. 5F) and BIM protein (BME_L_, BIM_L_, BIM_S_) (Fig. 5I, compare lanes 2, 3 and 4). Similar response to drug treatments were observed in the case of transcript levels of other p73 targets *CDKN1A* (encodes p21) and *BBC3* (encodes PUMA) in cells expressing mutant 53 (Fig.5G). Consistent with these effects on expression of p73 target genes, the combination treatment was more effective than mono treatments in enhancing apoptosis as determined by Annexin V assay (Fig. 5H, Supplementary Fig. S5A) and PARP cleavage assay (Fig. 5I) in a p73-dependent manner (Fig. 5H and 5I). The necessity of the expression of both mutant p53 and ERβ for the TNBC cells to be differentially responsive to Doxo monotreatment versus Tam+Doxo combination treatment is further supported by the same levels of TP73 RNA expression (Fig. 5C) and BIM protein expression (Fig. 5J) in cells depleted of both ERβ and mutant p53 (ERβ^KD^/p53KO) and low, but same levels of TP73 RNA (Fig. 5D) BIM protein levels (Fig. 5J) in cells where ERβ alone was depleted while mutant p53 was kept intact (ERβ^KD^/p53R280K). Apoptotic response as determined by Annexin V (compare Fig. 5H and 5K, Supplementary Figs. S5A, S5B), PARP cleavage (compare Fig. 5I and 5L), and BIM expression (compare Fig. 5I and 5L) assays was similar in MDA-231/LM2-4^Luc^ and MDA-MB-468 TNBC cells expressing different endogenous mutant p53 proteins.

Altogether, these data demonstrate the importance of ERβ-mutant p53-p73 axis in the response of TNBC cells to Tam + Doxo combination therapy

### Tamoxifen and Doxorubicin Reprogram TNBC Transcriptome to Regulate Multiple Pathways Converging on Cell Death by Apoptosis and Ferroptosis

To investigate mechanisms underlying the synergy between tamoxifen and doxorubicin in suppressing TNBC, we performed RNA-Seq to analyze differential gene expression (DGE) in MDA/231 LM2-4^luc^ cells in response to treatment with Tam and Doxo as single agents or in combination. Venn diagrams of DGE showed that in addition to the genes responsive to either of the two monotherapies, there is a unique set of genes that are responsive to the combination therapy (Suppl Fig. 6A). Gene set enrichment analysis (GSEA) of RNA-seq data revealed major cellular pathways impacted by monotherapies as well as combination therapy (Fig. 6A-H). In most of the cases, the pathways are affected in the same direction by either Tam or Doxo monotherapies with a much larger effect in response to the combination therapy. In the case of some gene sets (such as ‘reactome regulation of cholesterol biosynthesis’, ‘reactome iron uptake and transport’, and ‘reactome caspase activation via death receptors’) Tam and Doxo monotherapies regulate the gene sets in opposite direction; however, combination of these two drugs resulted in synergistic effect of upregulating tumor suppressive pathways.

**Figure 6.**
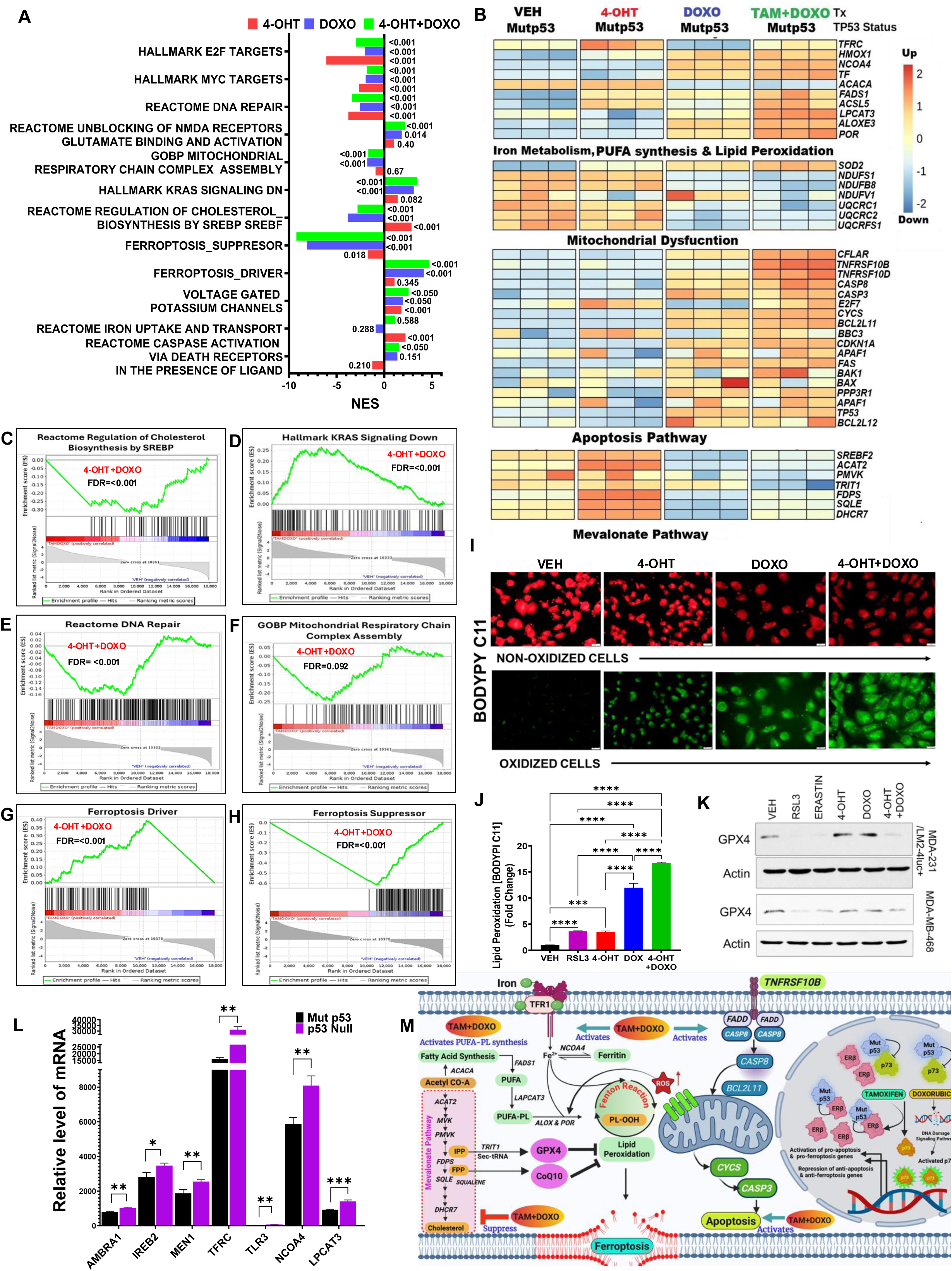
4-Hyrdoxy tamoxifen and doxorubicin combination reprograms TNBC transcriptome to regulate multiple pathways converging on cell death by ferroptosis and apoptosis. A, RNA-seq analysis of MDA-231/LM2-4 cells following treatment with Veh, 4-OHT (10 µM), DOXO (0.5 µM), or the combination of 4-OHT and DOXO for 24 h. Gene Set Enrichment Analysis (GSEA) was performed using the Molecular Signatures Database (MSigDB) C2 curated gene set collection to identify significantly enriched biological pathways. The plot displays the top enriched gene sets associated with each treatment condition, including both positive and negatively enriched pathways. Pathways enriched in 4-OHT-treated cells are indicated in red, those enriched in DOXO-treated cells in blue, and those enriched in the 4-OHT + DOXO combination group in green. Gene sets presented met the significance threshold of a normalized enrichment score (NES) with a false discovery rate (FDR; q-value) < 0.05 in the combination treatment group. **B,** Heatmaps showing the expression profiles of differentially expressed genes (DEGs) identified by RNA-seq analysis of MDA-231/LM2-4 cells treated as discussed in **A.** Genes are grouped according to their involvement in key biological pathways associated with ferroptosis and apoptosis response, including iron metabolism, polyunsaturated fatty acid (PUFA) synthesis and lipid peroxidation, mitochondrial dysfunction, apoptosis, and the mevalonate pathway. Heatmap color intensity represents relative gene expression levels upregulated and down regulated across treatment groups. **C-H,** Representative GSEA plots highlighting gene sets related to Cholesterol Biosynthesis by SREBP (**C**), KRAS signaling down (**D**), Reactome DNA Repair (**E**), and Mitochondrial Respiratory Chain Complex Assembly (**F**), Ferroptosis Driver (**G**) and Suppressor (**H)**. **I**, Representative fluorescence micrographs of BODIPY581/591C11 stained MDA-231/LM2- 4^LUC⁺^ cells showing (oxidized cells-green; non-oxidized cells- red) following treatment of Veh, 4- OHT (10 μM), DOXO (0.5 μM), or the combination for 24 h, followed by BODIPY581/591C11 staining (2 μM, 1 hour). **J**, Quantification of BODIPY581/591 stained cell on treatment as discussed in **(I).** Data represents the mean fold change of green/red fluorescence ratio. **K**, Immunoblot analysis of GPX4 expression in MDA-MB-468 and MDA231/LM2-4 cells treated with Veh, RSL3 (0.25 μM), Erastin (0.5 μM) (4-OHT (10 μM), DOXO (0.5 μM), or the combination of 4- OHT+ DOXO for 24 h. **L,** Basal level of ferroptosis driver gene expression in MDA231/LM2-4 vs p53KO from RNA-seq analysis of and following 24 h treatment discussed in **A. M,** Schematic illustration showing the proposed mechanism by which 4-OHT and DOXO synergistically induce ferroptosis and apoptosis in TNBC cells. RNA-seq data was analyzed using set of three independent experiments. Statistical analysis in Fig. **J** was assessed using one-way ANOVA and Fig. **L** using Student’s *t*-test. Statistical significance is indicated as follows: ns: non-significant, *P ≤ 0.05, **P ≤ 0.01, ***P ≤ 0.001, ****P ≤ 0.0001 significant.

Multiple genes in pathways that contribute toward regulation of cell death ferroptosis were differentially expressed in response 4-OHT + Doxo combination treatment. Expression of transferrin receptor C gene (*TFRC)* that encodes membrane transferrin receptor TFR1 involved in iron uptake; transferrin encoding gene (*TF*); and nuclear receptor coactivator 4 (*NCOA4*), a cargo receptor for ferritinophagy, were upregulated by both 4-OHT and 4-OHT + Doxo combination (Fig. 6B).

Acetyl CoA-carboxylase (*ACACA*), a repressor of ferroptosis, was downregulated by 4-OHT + Doxo combination. Genese regulating poly unsaturated fatty acid (PUFA) synthesis were downregulated by 4-OHT, and moderately upregulated by Doxo. However, combination of Tam+Doxo robustly increased expression of most of these genes (Fig. 3B). Examples include fatty acid desaturase 1 (*FADS1*) responsible for PUFA synthesis from MUFA and making TNBC cells more sensitive to ferroptosis (72), ACL5 (73), and *LAPCAT3*, an enzyme that modulates ferroptosis by regulating esterification of PUFA-CoA to form PUFA-phosphatidyl ethanolamine (PUFA-PL) (74), which, in turn, is oxidized to form PUFA-OOH. Production of ROS through iron-dependent Fenton reaction and activation of iron-containing enzymes such as lipoxygenases contribute toward this oxidation. p53 transcriptional target SAT1 (75) and downstream lipoxygenase *ALOXE3,* an activator of PUFA synthesis and enhancer of ferroptosis are upregulated in response to combination treatment. NADPH-cytochrome p450 reductase (*POR*) (76) are also upregulated.

Mitochondria is another major source of ROS. Electrons leaking from defective electron transport chain (ETC) complexes I and III produces superoxide, which is converted to hydrogen peroxide (H_2_O_2_) by superoxide dismutase (SOD). H_2_O_2_ reacts with Fe2^+^ leading to generation of hydroxyl radicals (*OH), then to PUFA radicals (PUFA*), PUFA peroxy radicals (PUFA-OO*), and ultimately to PUFA hydroperoxide (PUFA-OOH), leading to ferroptosis (77). Importantly, 4-OHT + Doxo combination downregulated ETC complex I subunits *NDUFS1*, *NDUFB8*, and *NDUFV1* along with ETC complex II subunits UQCRC1, *UQCRC2*, and *UQCRSF1*, whereas *SOD2* was upregulated by the drug combination treatment. GSEA plot revealed mitochondrial respiratory chain complex assembly pathway down regulated by the drug combination (Fig. 6B, 6F; Supplementary Figs. S6P, S6Q).

The mevalonate pathway is responsible for synthesis of cholesterol and non-steroidal isoprenoids. Several mevalonate pathway enzymes are regulated in opposite direction (upregulation by 4-OHT; downregulated by Doxo) by mono treatments, whereas these enzymes were downregulated by Tam + Doxo combination treatment (Fig. 6A, 6B, 6C, and Supplementary Figs. S6L, S6M). SREBP2 (encoded by *SREBF2* gene), the master transcriptional regulator of this pathway, is blocked by WTp53-mediated induction of the ABCAI cholesterol transporter gene. Lack of this inhibition in mutant p53-expressing cells along with direct transcriptional activation of multiple mevalonate pathway genes by interaction of mutant p53-SREBP2 protein complex with the gene promoter regions, leads to upregulation of mevalonate pathway (12,78). Although SREBF2 expression is upregulated by 4-OHT, 4-OHT + Doxo combination drastically decreases it in the MBA231/LM2-4 cell line. Consistent with this, GSEA plot showed downregulation of cholesterol biosynthesis pathway by SREBP (Fig. 6C). ACAT2, a cytoplasmic multifunctional enzyme that catalyzes two molecules of acetyl-CoA to form an acetoacetyl-CoA and a CoA, plays a crucial role in lipid metabolism including fatty acid β-oxidation and de novo cholesterol synthesis. Additionally, ACAT2 activates GCLM by lactylation resulting in increased GSH and inhibition of ferroptosis (79). *ACAT2* expression is upregulated by Tam, whereas it is downregulated by Tam+ Doxo combination. Furthermore, mevalonate diphosphate decarboxylase (*MVD*) that synthesizes isopentyl 5-diphosphate (IPP), and farnesyl diphosphate synthase (*FDPS*) that synthesizes farnesyl diphosphate (FPP) are upregulated by 4-OHT, while both are downregulated by Doxo as well as 4-OHT + Doxo combination. IPP, a substrate for tRNA isopentyl transferase 1 (*TRIT1*) is necessary for the maturation of selenocysteine tRNA (Sec-tRNA) required for translation of GPX4, a major antioxidant that inhibits lipid peroxidation and ferroptosis (80). *TRIT1* is downregulated in response to combination treatment. Clinical relevance of GPX4 levels in TNBC tumors was reported recently (81). Farnesyl-PP-Synthase (*FDPS*) that synthesize farnesyl-PP (FPP), a precursor for coenzyme Q10 (CoQ10) which functions as a lipophilic radical-trapping anti-oxidant that inhibits propagation of lipid peroxides and ferroptosis (82) is repressed by Tam + Doxo combination. Ferroptosis suppressor protein 1 (FSP1) plays an important role in CoQ10 function. Ubiquinol (reduced form of CoQ10) traps lipid peroxyl radicals that mediate lipid peroxidation, whereas FSP1 catalyzes the regeneration of CoQ10 using NAD(P)H (83,84). Another target of repression by Tam + Doxo combination is squalene epoxidase or squalene monooxygenase (encoded by *SQLE*), the second rate-limiting cholesterol biosynthesis enzyme downstream of *HMGCR*, converts nonsterol intermediate squalene to 2,3(S)-oxidosqualene. Squalene has anti-ferroptosis activities and it accumulates in cells devoid of squalene monooxygenase making the cells resistant to ferroptosis (85).

Of note, genes involved in downregulation of KRAS signaling were upregulated in response to Tam+Doxo combination treatment (Fig. 6D). MDA-MB-231 cells carry mutations in KRAS and RAS effector pathways (86) and oncogenic KRAS reshapes cancer cell metabolism and activate pathways that confer resistance to ferroptosis. Several MYC targets including ferroptosis suppressor *SREBF2* are downregulated (Fig. 6A, 6B, and Supplementary Fig. S6C). Importantly, GSEA analysis also revealed downregulation of DNA damage repair pathways (Fig. 6 E). Downregulation of DNA damage response (DDR) can activate ferroptosis (87) and is consistent with upregulation of E2F7 (Fig. 6B), a contributor toward attenuation of DNA repair function through transcriptional repression of genes including *RAD51* (60).

Overall, as shown in the GSEA plots (Fig. 6G and 6H; Supplementary Figs. S6N, S6O), RNA-seq data show that 4-OHT+ Doxo combination upregulates pro-ferroptosis pathways and down regulates ferroptosis suppressor pathways. Pro-ferroptosis PUFA synthesis and oxidation coupled with increased iron uptake and release of Fe2+ causing increased Fenton reaction and ROS production. Functional mitochondrial destabilization leads to further increase in ROS contributing toward increased lipid peroxidation and ferroptosis. This pro-ferroptosis effect of the drug combination is complemented by suppression of expression of multiple genes encoding various enzymes in the mevalonate pathway leading to decrease in levels of GPX4 and CoQ10, two major antioxidants that inhibit ferroptosis (Fig. 6A-C; Supplementary Figs. S6L, S6M, S6Q). The RNA-seq data is corroborated by fluorescent lipid peroxidation-sensitive dye BODIPY581/591-C11 oxidation data in MDA/231-LM2-4^luc^ and MDA-MB-468 TNBC cells in response to 4-OHT, Doxo, and 4-OHT + Doxo treatment (Fig. 6I and 6J; Supplementary Fig. S6V, S6W). Furthermore, immunoblot of protein extract from treated cells showed that 4-OHT + Doxo combination results in reduction of GPX4 levels in both these cell lines Fig. 6K).

4-OHT + Doxo combination treatment led to reprogramming of gene expression that was not only pro-ferroptosis, but also pro-apoptosis Fig. 3D, and Fig. 6B and 6C). Consistent with upregulation of apoptosis in response to the drug combination as was evident from increased cleaved PARP, cleaved caspase 3 (Fig. 3G), Annexin V+ assay (Fig. 3I; Fig. 4F-4J), RNA-Seq data showed increased expression of several pro-apoptosis genes (Fig. 3D, Fig. 6B and 6C).

Importantly, most of the differentially regulated (upregulated as well as downregulated) ferroptosis and apoptosis regulatory pathway genes showed dependency on the context of mutant p53 expression. For example, basal expression levels of ferroptosis activators *BCL211*, *IREB2, MEN1, SOS1, TFRC, TLR3, NCOA4,* and *LPCAT3* were lower whereas ferroptosis suppressors *STC1, PIM1, SLC2A12* were higher in cells expressing mutant p53 (MDA-231LM2-4) compared to isogenic cells without p53 (MDA-231LM2-4p53KO) (Fig. 6L and Supplementary Table S1) demonstrating the ability of mutant p53 to suppress ferroptosis. Based on these data, activation and repression of expression of multiple genes regulating pathways of extrinsic and intrinsic cell death, cholesterol synthesis, iron metabolism, PUFA synthesis, and lipid peroxidation, and mitochondrial function converge on increased apoptosis and ferroptosis as shown schematically in the working model (Fig. 6M).

### Tamoxifen and Doxorubicin Combination is Effective in Suppressing TNBC Cell Line-Derived Xenograft (CDX) and Patient-Derived Xenograft (PDX) Tumors

Next, we analyzed if our observations in TNBC cell line models can be recapitulated in *in vivo* TNBC models. We generated tumor cell line-derive xenografts (CDXs) in SCID mice from isogenic MDA231/LM2-4^Luc^ human TNBC cells expressing p53 with different mutations (R280K and R248Q), or cells with wild type p53, or p53 knockout (p53KO) cells, and cells where ERβ was stably knocked down. The mice were treated with vehicle, Tam, Doxo, or Doxo + Tam combination. PK/PD analysis (Supplementary Figs. S7A-S7E) showed that Tam synergized with Doxo in inhibiting growth of tumors generated from isogenic MDA/231/LM2-4^Luc^ cells expressing either mutant p53 R80K (Fig. 7A) or mutant p53 R248Q (Supplementary Fig. S7F) suggesting combination therapy was much more effective than either Tam or Dox monotherapies in suppressing growth of xenograft tumor with mutant p53. The synergistic effect of the drugs was not observed in the case of xenograft tumors derived from isogenic cells with p53 knockout (MDA/231^Luc^ p53KO) (Fig. 7B) or in those expressing wild type p53 (MDA/231^Luc^ WTp53KO) ((Fig. 7C). Requirement of ERβ for tumor growth and response to drugs was assessed in MDA/231^Luc^ cells where ERβ was depleted (Fig. 7D). The tumors generated from these cells were not responsive to Tam + Doxo combination suggesting the necessity of ERβ for the response. As the isogenic cell lines share the same genetic characteristics except for the difference in p53 or ERβ status, positive therapeutic response to Tam + Doxo can be ascribed directly to the presence of both mutant p53 and ERβ. Consistent with our data from *in vitro* TNBC cell line model systems, ERβ-mutant p53 interaction in tumors was increased in response to the combination therapy (Fig. 7E), whereas expression of proliferation marker ki67 was decreased (Fig. 7F) and ERβ protein levels were increased (Supplementary Fig. S7H). Body weight profile during treatment showed lack of toxicity of the treatment regimens (Supplementary Fig. S7G).

**Figure 7.**
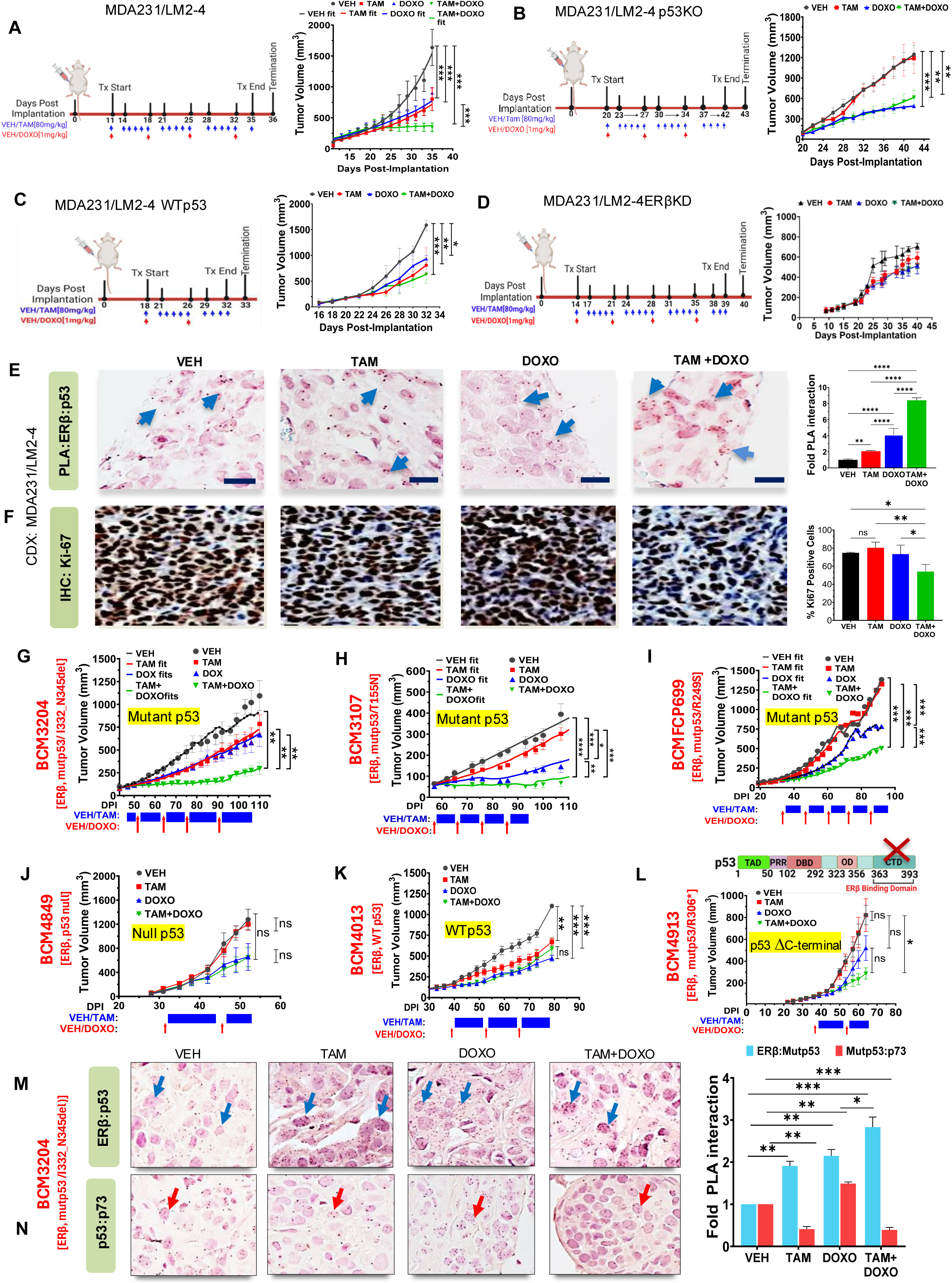
Tamoxifen and doxorubicin combination suppresses tumor growth *in vivo*. A–D, Treatment schema for cell-derived xenograft (CDX) models and corresponding tumor growth analysis. SCID mice bearing MDA-MB-231/LM2-4 tumors generated from p53R280K (**A**), p53KO **(B)**, WTp53 **(C),** or ERβKD **(D)** were treated with vehicle (corn oil and PBS), Tamoxifen citrate (TAM; 80 mg/kg, oral gavage), or doxorubicin (DOXO; 1 mg/kg, intraperitoneal injection) and tumor volumes were measured at the indicated time points. Data are presented as mean ± SEM (n = 5 mice per group). Statistical significance was determined by one-way ANOVA. In Fig. **A,** Tumor volume plot: Symbols (●,▪,▴&▾) represent observed tumor volume data, and lines represent the model-fitted values. **E**, Representative Brightfield micrographs showing ERβ-Mutant p53 interaction in tumor tissue following treatment assessed by Brightfield PLA (scale bar = 100 µm). Right panel: quantified PLA dots of 300 cells per treatment. **F,** Representative images of Ki67 IHC across all treatment groups (Scale bar 100 μm). Right panel: Ki67 expression H-Score was measured and quantified using GraphPad Prism 9. **G–L**, Treatment schema and tumor growth analysis of patient-derived xenograft (PDX) models. PDXs with high mutant p53 expression BCM3204 **(G)**, BCM3107 **(H)**, BCMFCP699 **(I)**, and BCM4849 p53-null (**J**), BCM4013 WTp53 (**K**), and BCM4913 C-terminally truncated p53 **(L)** models were treated with vehicle (corn oil and PBS), TAM; 80 mg/kg, oral gavage), or doxorubicin (DOXO; 1 mg/kg, intraperitoneal injection) and tumor volumes were measured at the indicated time points. Data are presented as mean ± SEM (n = 5 mice per group). Statistical significance was determined by ANOVA. **G–I,** In tumor volume plot: Symbols (●,▪,▴&▾) represent observed tumor volume data, and lines represent the model-fitted values. **M–N,** Representative Brightfield PLA micrographs showing ERβ**–**Mutant p53 interaction (**M**) and p53–p73 interaction (**N**) in PDX tissue BCM3204, (scale bar; 100 µm). Right panel; quantified PLA dots of 300 cells per treatment. Quantitative data in **A-N** were plotted and analyzed in GraphPad Prism 9. Error bars denote ± SEM. Statistical analysis was performed using one-way ANOVA. Statistical significance is indicated as follows: ns: non-significant, *P ≤ 0.05, **P ≤ 0.01, ***P ≤ 0.001, ****P ≤ 0.0001 significant.

For addressing relevance of our data from cell lines and CDX models to TNBC patient tumors, we used well-characterized patient-derived xenograft (PDX) models (88). Expression levels of ERβ and p53 proteins in the patient tumor tissues used for implantation along are shown (Supplementary Fig. 7I) along with age at diagnosis, p53 status of tumor tissues, and race (Supplementary Fig. 7J). As in the case of CDXs, PDXs expressing mutant p53 (I332-N345 deletion of the oligomerization domain), T155N, R249S (Fig. 7G, 7H, and 7I, respectively), and R248Q (Supplementary Fig. S7K) responded to the combination therapy better than monotherapies with either Tam or Doxo, whereas PDX tumor that is p53 null (Fig. 7J) and the one expressing wild type p53 (Fig. 7K) did not show synergistic response. Importantly, PDX 4913 with termination codon at aa 306 resulting in p53 devoid of C-terminal domain (CTD) that is necessary for interaction with ERβ (37,44) did not show synergistic response (Fig. 7L). These data suggest that mutant p53-ERβ interaction is required for synergistic effect of Tam + Doxo combination therapy *in vivo* in human TNBC tumors. Predictive distribution for the PDX study combination model and the Goodness of fit model performance in the combination therapy observed versus individual model predicted values demonstrated synergism of the two drugs in the combination therapy (Fig. 7G-I and Supplementary Fig. 7A-7E). Both ERβ and mutant p53 were expressed in the tumor tissues (Supplementary Figs. S7M, S7N) and PLA revealed that ERβ-mutant p53 interaction was increased (Fig. 7M) in response to the combination therapy while mutant p53-p73 interaction was decreased (Fig. 7N). Body weight profile during treatment showed lack of toxicity of the treatment regimens (Supplementary Fig. S7L). These data demonstrate that Tam + Doxo combination effectively suppresses growth of TNBC CDX and PDX tumors expressing mutant p53 and ERβ.

## DISCUSSION

Tam has not been used for TNBC therapy as ERα, the canonical target of Tam, is not expressed in this breast cancer subtype. This study, using pharmacological and genetic approaches in TNBC cell line models, and tumor cell line-derived and patient-derived xenograft (CDX and PDX) models has challenged this conventional paradigm and show that Tam in combination with Doxo has the potential to be repurposed as a new therapeutic strategy against TNBC.

Based on reports of tumor suppressive role of ERβ in TNBC (26), it is logical to assume that 17β- estradiol (agonist for both ERα and ERβ) as well as ERβ-specific agonists such as DPN and LY500307 may exert tumor suppressive activity. However, although pre-clinical studies in mouse xenograft tumor models have shown tumor-suppressive activities of these agonists (89), a phase 2 clinical study of high-dose estradiol in metastatic TNBC (NCT01083641) was closed as a result of limited efficacy (90). Our finding that Tam, but not PHTPP (ERβ antagonist) or 17β-estradiol or DPN (ERβ agonists) were able to increase ERβ-mutant p53 interaction and mediate blockage of tumor cell proliferation and increase cell death (Figs. 1 and 2; Supplementary Fig. S1) suggest that anti-TNBC effects of Tam cannot be explained by traditional paradigms of receptor agonists band antagonists. Agonism and antagonism of steroid hormone receptors have been historically defined based on their ability to access their cognate binding sites on their target gene promoters and regulate transcription of these genes. Based on this definition, Tam is considered as a SERM that acts as an antagonist of estrogen receptors in the breast tissue. However, our data show that the effect of Tam in breast tumors is not confined to this canonical paradigm. Our data show that Tam impacts interaction of ERα and ERβ with p53. In a window-of-opportunity clinical study in ERα-positive breast cancer patients, we have shown that Tam disrupts ERα-Wtp53 protein-protein interaction in tumors resulting in reactivation of p53 leading to tumor suppressive transcriptomic reprogramming (44,91). In the case of TNBC, Tam stabilizes ERβ and increases ERβ-mutant p53 interaction resulting in reactivation of p73 leading to tumor suppressive effects in cell lines (37) and *in vivo* in human tumors (39,44). RNA-seq analysis showed that Tam upregulated BTG2, E2F7, BBC3, TIMP2 while repressing CXCL1 and LSM4 in the MDA- 231/LM2-4 cells expressing mutant p53, whereas the effect was reversed in MDA-231/LM2-4 p53KO cells suggesting vulnerability of cells expressing mutant p53 to anti-tumorigenic effects of Tam. Importantly, pharmacodynamic studies revealed that Tam when combined with Doxo resulting in synergy while the synergy was lost in isogenic cells where mutant p53 was knocked out suggesting that an underlying cause of synergy is inhibition of oncogenic gain-of-function of mutant p53. Synergy occurs when the overall effect of the drug combination is greater than additive effects of each individual agent in the combination (92). Analysis of proliferation response showed that in all isogenic TNBC cell lines with different p53 mutations, IC50 for 4-OHT + Doxo was lower than that for Tam alone (Fig. 4C; Supplementary Fig. S6C) suggesting Tam synergized with Doxo to suppress proliferation. That in cells where mutant p53 was knocked out, there was no difference in IC50 for 4-OHT versus 4OHT + Doxo further strengthens the necessity of mutant p53 for cells being sensitive to the combination treatment. Of note, IC50 for combination therapy was higher than that of monotherapy with Tam in cells with WTp53. These observations were reinforced by synergy of Tam with Doxo in TNBC CDX and PDX tumors expressing mutant p53, but not in tumors with WTp53 or those with no p53. Furthermore, cells with depletion of ERβ did not respond to the combination therapy. Importance of ERβ for the synergistic effect of Tam + Doxo was further ascertained by the observation that a PDX model derived from a TNBC tumor with a deletion of the C-terminal domain (CTD) of p53 that we had reported to be necessary for binding to ERβ (37), was not responsive to the combination therapy (Fig. 7L). The unstructured CTD participates in every aspect of p53 functioning (93). It is a target of large number of post-translational modifications that may vary depending on cell and tissue type and nature of stress signaling (94,95). Many CTD-driven interactions are low affinity potentially ensuring rapid and flexible functional control. Topoisomerase II, a target of Doxo, has been reported to bind to the C-terminal of p53 (96). It is tempting to speculate that ERβ binding to the C-terminus may enhance susceptibility of topoisomerase II poisoning by Doxo, thus making the tumor with intact mutant p53 responsive to the combination therapy, while making the tumors expressing p53 devoid of CTD nonresponsive. Although ERβ-independent roles of CTD cannot be ruled out, our data on higher IC50 for Tam + Doxo combination upon depletion of ERβ in TNBC cells expressing mutant p53 with intact CTD, along with non-responsiveness of CDX tumors depleted of ERβ to the drug combination, argues in favor of the necessity of ERβ binding to p53 CTD. In vitro and in vivo data from the current study collectively demonstrate that both ERβ and mutant p53 are required for sensitizing the tumors to Tam + Doxo therapy.

Multiple mechanisms underlie the synergy between Tam and Doxo. Our results demonstrate efficient interaction between ERβ and mutant p53 while decreasing p73-mutant p53 interaction leading to reactivation of p73 and cell death in TNBC cell lines and tumors in response to Tam + Doxo combination therapy. Consistent with this finding, protein depletion by RNA knockdown or gene deletion has revealed the requirement of ERβ, mutant p53, and p73 for differentially increasing death in response to Tam +Doxo combination as compared to monotherapies by either Tam or Doxo. Analysis of differential gene expression in cells exposed to either monotherapies or combination therapy revealed that multiple genes in various pathways converging on apoptosis and ferroptosis preferentially activated in response to combination therapy as compared to monotherapies. Several pathways are affected in the same direction by either Tam or Doxo monotherapies with a much larger effect in response to the combination therapy. For example, iron metabolism, PUFA synthesis and peroxidation, and apoptosis pathway genes are modestly activated by either Tam or Doxo while the activation is robust in response to the combination of these two drugs. On the other hand, several genes encoding mitochondrial energy metabolism genes are modestly downregulated by monotherapies while the downregulation is robust in response to combination therapy potentially leading to mitochondrial dysfunction and ferroptosis. Of note, in the case of mevalonate pathway, several genes involved are upregulated by Tam where as Doxo downregulates them. In response to the drug combination, the pathway genes are downregulated, potentially leading to depletion of major antioxidants GPX4 and Coenzyme Q10 that are known inhibitors of ferroptosis. That the genes responsive to Tam + Doxo combination therapy converge on apoptosis and ferroptosis are corroborated by the data from our apoptosis and ferroptosis assays.

Our findings have important clinical implications. Chemotherapeutic agents such as Doxo are integral to TNBC therapeutic regimens such as targeted ADCs as well as immune therapies including immune checkpoint inhibitors (ICIs). However, adverse effects such as cardiotoxicity from cumulative doses of anthracyclines such as Doxo has been a major clinical challenge. Whie Tam as a monotherapy is not effective against TNBC, our finding that Tam when repurposed in combination with Doxo lowers the IC50 of Doxo is important from the perspective of reducing adverse effects. Our data suggest Tam enhances ERβ-mediated activation of p73 and down regulates DNA damage repair pathway genes while DNA damage is increased by Doxo. Combination of these processes effectively enhances cell death by both apoptosis and ferroptosis. Therapy-resistant state of cancer is dependent on lipid peroxidase pathway (51) and this vulnerability can be exploited to induce ferroptosis by Tam + Doxo combination. Spatial, temporal and mechanistic regulation of cell death is critical for the body to defend against illnesses including cancer. Recent studies in cancer cell lines and in vivo in mouse models have revealed important roles for mutant p53 in suppressing ferroptosis and apoptosis in cancer (9,52–54). Cancer stem cell-regulated phenotypic plasticity has also been reported to protect metastasized cancer cells from ferroptosis (97). Robust induction of ferroptosis in addition to apoptosis response to the combination therapy could kill the tumor cells that have become resistant to apoptosis, and this could be a major factor contributing to synergy. Importantly, majority of apoptosis and ferroptosis genes respond to the combination therapy in a mutant p53-dependent manner (Table S2). Therefore, this novel drug repurposing approach has the potential to be very effective to exploit the mutant p53 vulnerability in more than 80% of TNBC.

## METHODS

### Cell Culture

The following cell lines were used in this study: MDA-MB-468 obtained from American Type Culture Collection (ATCC) and MDA231/LM2-4^LUC+^ (provided by Dr. John Ebos, RPCCC) (57). MDA231/LM2-4^LUC+^ p53KO line was generated from MDA231/LM2-4^LUC+^ as described previously (37). Stable isogenic cell lines were generated from MDA231/LM2-4^LUC+^ p53KO, including p53 WT, p53 mutant variants (R175H, R273H, R248Q, R280K, and wild-type p53), ERβ knockdown (ERβKD/p53KO), and ERβ knockdown/mutant p53 R280K (ERβKD/p53R280K) (see Supplementary Fig. S4A). All cell lines were cultured in Dulbecco’s Modified Eagle’s Medium (DMEM) (Corning, Corning, NY, USA, Cat #10-013-CM), supplemented with 10% fetal bovine serum (FBS) (GIBCO/ThermoFisher, Waltham, MA, USA, Cat #20937-028), and penicillin-streptomycin (50 µg/mL) (Corning, NY, USA, Cat #30-002-CI). Cultures were maintained at 37°C in a humidified incubator with 5% CO₂. Mycoplasma testing was performed using a PCR-based detection kit in the shared facility of RPCCC.

### Generation of stable Isogenic TNBC cells with different p53 and ERβ status

Cloning of Wild-Type and Mutant TP53 via Restriction Digestion: Stable MDA231/LM2-4LUC+ p53KO cell lines constitutively expressing wild-type p53 or mutant p53 variants (R175H, R273H, R248Q, and R280K) were generated using the lentiviral vector TLO1001.CV (Cat. #15536), obtained from TransOMIC Technologies Inc. Restriction enzyme mediated cloning was used to clone p53WT, mutant p53 (R175H), and mutant p53 (R273H) in lentiviral vector TLO1001.CV. Briefly, cDNA encoding p53 WT and mutant p53 (R175, and R273H) was PCR amplified from pCR3.1 vector using the following primers: TP53 forward 5’- TGGTGGAATTCATGGAGGAGCCGCAGTCAGATCC-3’ and TP53 reverse 5’- CGCGGGATCCTCAGTCTGAGTCAGGCCCTTCTGT-3’. The TLO1001.CV vector and the p53 cDNA insert were digested overnight at 37°C with the restriction enzymes EcoRI and BamHI. Following digestion, both the vector and insert were column-purified and subsequently ligated using the Quick Ligation™ Kit (NEB, Cat. #M2200S) for 10 minutes at room temperature. The resulting ligation products were transformed into competent *Escherichia coli* K802 cells, generously provided by Dr. Gerald Koudelka, and selected on ampicillin-containing agar plates. The positive clones were confirmed by Sanger sequencing.

Generation of TP53 Mutants via Site-Directed Mutagenesis: The Tlo1001.CV TP53 WT vector was mutagenized into p53R248Q and p53R280K using the following primers. R248Q (FOR: CATGAACCAGAGGCCCATCCTCACCATC, REV: GGCCTCTGGTTCATGCCGCCCATGC), p53R280K (FOR: TCCTGGGAAAGACCGGCGCACAGAGGAA, REV: CGGTCTTTCCCAGGACAGGCACAAACAC) Briefly, the Tlo1001.CV TP53 WT vector was PCR amplified using site-directed mutated primers with High-Fidelity DNA polymerase Phusion (New England Biolabs, Ipswich, MA, USA, Cat# M0493S). The thermocycler program: an initial denaturation at 95 °C for 5 minutes, followed by 20 cycles of 95 °C for 1 minute, 65 °C for 30 seconds, and 72 °C for 8 minutes, with a final elongation at 72 °C for 10 minutes. The PCR products were subsequently digested with DpnI endonuclease (New England Biolabs, Ipswich, MA, USA, Cat# R0176S) at 37 °C, overnight. DNA was column-purified and transformed into competent *Escherichia coli* K802 and selected on ampicillin-containing agar plates. The positive clones were confirmed by Sanger sequencing.

Generation of ERβ knockdown construct: The miR-30 system was employed to design multiple shRNAs specifically directed against the *ESR2* transcript. The PLV ESR2KD vector (Vector ID: VB230220-1180pye), which constitutively expresses shRNAs targeting *ESR2*, was obtained from Vector Builder. The target sequences for *ESR2* are as follows: *ESR2* transcript location Exon2 (target seq- TTAGTGGTCCATCGCCAGTTAT, TAGACAGCCACCATGAATATCC, and ATGAATTACAGCATTCCCAGCA) Exon3 (target seq- AGGGATGCTCACTTCTGCGCTG).

Transduction into human cells and clonal selection: Lentiviral particles were produced in HEK 293T cells. Briefly, psPAX2 (packaging vector, 2.5 µg), pMD2.G (envelope vector, 1.0 µg), and Tlo1001.CV vector carrying p53WT, mutant p53 (R175H, R248Q, R273H, R280K), or PLV ESR2 KD vector (2.5 µg) were transfected using LipoD293 Reagent (SignaGen Laboratories, Cat # SL100668) following the manufacturer’s guidelines. The culture supernatant containing viral particles was harvested at 48 and 72 h and passed through a 0.45 µm cellulose acetate syringe filter. MDA231/LM2-4^LUC+^ p53KO cells were transduced with the filtered viral supernatant for 24 h. The cells were selected with 6 µg/ml blasticidin for Tlo 1001.CV vector and 2 µg/ml puromycin for PLV ESR2 KD vector. Single clones were isolated according to Addgene’s protocol for “Isolating a Monoclonal Cell Population by Limiting Dilution” (Addgene protocol: https://www.addgene.org/protocols/limiting-dilution).

### Ligands and Drugs

DPN (Cat#1494), PHTPP (Cat#2662), 4-OH-tamoxifen (Cat#3421), Tamoxifen Citrate (Cat#0999), Doxorubicin Hydrochloride (Cat#2252), and β-Estradiol (Cat#2824), were sourced from TOCRIS, Bio-Techne Corporation, MN, USA. Ferroptosis inducer Erastin (Cat#HY-15763) and RSL3 (Cat#HY-100218A) were purchased from MedChemExpressLLC, NJ, USA.

### siRNAs Transfection and Knockdown

Gene silencing experiments were performed by transiently transfecting cells with non-specific stealth control siRNA (Invitrogen/Thermo Fisher, Waltham, MA, USA, Cat #12935-300), 120 nM of ESR2 (Cat# HSS103380; sequence: 5’-CCCUGCUGUGAUGAAUUACAGCA-3’), and 100 nM of TP73 (Cat# AM16708; sequence: 5’-CCAGACAGCACCUACUUCGtt-3’) for 48h. A non-targeting Stealth siRNA (ThermoFisher Scientific, Cat# 4390844,) was used as a negative control.

Lipofectamine 3000 (Thermo Fisher Scientific, Waltham, MA, USA, Cat #L3000015) was used for transfection following the manufacturer’s protocol.

### RNA Isolation and Quantitative real-time polymerase chain reaction (qRT-PCR)

RNA was extracted using TRIzol reagent (Invitrogen, Waltham, MA, USA, Cat #15596026) using the manufacturer’s instructions. The purified RNA was quantified using Nanodrop 8000 spectrophotometer (Thermo Fisher, Cat #ND-8000-GL), digested with DNase I (amplification grade I (Thermo Scientific, Cat #EN0521)), and cDNA was synthesized using cDNA Synthesis Kit (Bio-Rad iScript, Cat # 1708890). qRT-qPCR was conducted using iTaq Universal SYBR Green Supermix (Bio-Rad, Hercules, CA, USA, Cat #1725121) following the standardized protocol (37,98) The relative gene expression was determined using the ΔΔCT method with β-actin as the reference gene.

### Oligonucleotides

For quantitative RT-PCR analyses, the following primer sequences (5′–3′) were used: *ESR2* (ERβ) forward, CCTATGTAGACAGCCACCATGAAT; reverse, CCCACCTCCCAAGTTAGTGACATT. β-Actin forward, ATGGGTCAGAAGGATTCCTATGT; reverse, AAGGTCTCAAACATGATCTGGG. *CDKN1A* (p21) forward, GAGACTCTCAGGGTCGAAAACG; reverse, GATGTAGAGCGGGCCTTTGA. *BBC3* (PUMA) forward, ATGCCTGCCTCACCTTCATC; reverse, TCACACGTCGCTCTCTCTAAACC. *BCL2L11* (BIM) forward, GGCCCCTACCTCCCTACA, reverse, GGGGTTTGTGTTGAATTTGTCA. *TNFRSF10B* forward, GTGATTCAGGTGAAGTGGAGC reverse, CGACCTTGACCATCCCTCTG, *CASP8* (Caspase 8) forward, CCAGAGACTCCAGGAAAACAG reverse, GATAGAGCATGACCCTGTAGGC, *GPX4*, forward, GTGAGGCAAGACCGAAGTAAA reverse, GAACTGGTTACACGGGAAGG *TP73* (p73) forward, GCACCACGTTTGAGCACCCT reverse, GCAGATTGAACTGGGCCAGA *TFRC* forward, TTTCCACCATCTCGGTCATC; reverse, GGGACAGTCTCCTTCCATATTC.

### Wound Healing Assay

For wound healing (scratch) assays, self-adhering biocompatible silicone Ibidi Culture-2 chambers (Product# 80209) were placed into 24-well plates using the manufacturer’s instructions. MDA231/LM2-4^LUC+^ and MDA231/LM2-4^LUC+^ p53KO cells were seeded at a density of 5.70 x 10^5^ cells/mL and cultured for 24 h. Upon reaching confluence, inserts were removed to generate a uniform 500 µm cell-free gap. Cells were treated with VEH, TAM (10 μM), DOXO (0.5 μM), and a combination of both, in triplicate, using the Tecan D300e Digital Dispenser. After treatment, the plate was placed in the BioTek Cytation 5, and images were acquired using a 4x objective set to phase contrast. The plate was then transferred to the BioTek BioSpa 8 Incubator for automated reading, with images taken from each well every hour for 24-48 h. The gap closure was monitored using Agilent BioTek Lionheart FX automated microscope (Santa Clara, CA, USA) and analyzed with Agilent BioTek Gen5 microplate reader and imager software. The analysis parameters were set to measure the confluency of each side of the wound as cells moved into the empty area. The percentage of confluency in the wound area was calculated and plotted using GraphPad Prism version 9.

### Matrigel Invasion Assay

Briefly, individual Corning PET membrane permeable supports (Corning #351152) were placed into Falcon 24-well companion plates (Corning #353504) to evaluate differences in invasion in response to treatment across multiple cell lines. The inserts were coated with Matrigel at a concentration of 250 μg/mL and set to polymerize at 37°C in 5% CO_2_. MDA231/LM2-4^LUC+^ and MDA231/LM2-4^LUC+^ p53KO cells were labeled with a fluorescent dye CellTracker™ Blue CMF2HC dye (ThermoFisher Scientific, Cat# C12881) at a concentration of (10 µM) to detect migration through the invasion membrane. Following Matrigel polymerization, cells were harvested, counted, and diluted to a concentration of 3.0 x 10^5^ cells in 5 mL of serum-free media. A 500 µL aliquot of each cell line suspension was added to the appropriate inserts. Treatments, including Veh, TAM (10 µM), DOXO (0.5 µM), and a combination of TAM and DOXO, were dispensed into the wells using a Tecan D300e Digital Dispenser. Subsequently, 750 µL of media containing 10% FBS was added below each insert to serve as a chemoattractant. Once the plates were fully set up, they were placed into the BioTek BioSpa8 robotic incubator and imaged using the BioTek Cytation5 multimode imaging system. CMF2HC blue fluorescence was captured automatically with the DAPI filter system at 4x magnification. Montage tiles of each well were taken every hour for 48 h. Image processing and analysis were performed using BioTek Gen5 Image Prime software.

### Immunoblotting

Cells were harvested by scraping into ice-cold 1x PBS and lysed in RIPA lysis buffer (50 mM Tris-HCl pH 8.0, 1% NP-40, 150 mM NaCl, 0.5% deoxycholate, 0.1% SDS, 5 mM EDTA) supplemented with EDTA-free “Roche cOmplete” protease inhibitor cocktail (Millipore Sigma, St. Louis, MO, USA, Cat #11873580001). Protein quantification was performed using Bradford reagent (BioRad, USA Cat #5000006). Equal amounts of protein were resolved by SDS-PAGE and transferred onto PVDF membranes. Membranes were blocked in 5% nonfat milk and incubated with primary antibodies overnight at 4 °C. HRP-conjugated secondary antibodies were used for detection by chemiluminescence. Antibodies and their respective dilutions used are provided in Table S1.

### Co-Immunoprecipitation Assay

Cells were harvested and lysed in ice-cold RIPA lysis buffer supplemented with a protease inhibitor cocktail (Millipore Sigma, Cat #11873580001). The cell lysates were then pre-cleared with Pierce™ Protein G Plus Agarose (Life Technologies/Thermo Fisher, Waltham, MA, USA, Cat #22852) and subjected to immunoprecipitation using specific antibodies against ERβ, p53, and IgG as a control, as previously described protocols (98). The protein complexes were eluted in 2× sample buffer and processed for immunoblotting as described above in Immunoblotting. The antibody details are provided in Supplementary Table S2.

### Cell Cycle Analysis

Following treatment, the cells were pulsed with BrdU (10 μM) for 1 hour. Cells were harvested, washed 2x with ice cold phosphate-buffered saline and fixed in 70% ice-cold ethanol. DNA was denatured using 2M HCl (hydrochloric acid). Cells were stained with BrdU Mouse mAB (BD Pharmingen, Cat #555627) overnight at 4 °C in dark. DNA was stained with Propidium Iodide (PI) (ThermoFisher, Cat #P1304MP) supplemented with PureLink™ RNase A (Invitrogen, Cat #12091039). The samples were then analyzed by flow cytometry. At least 20,000 events were collected per sample using BD LSRFortessa flow cytometer and analyzed using FCS Express 7. The results are shown as mean values ± standard deviation, based on three independent experiments.

### Cellular Senescence Assay

Cellular Senescence was assessed by staining with β-galactosidase (SA-β-gal) staining kit (Millipore Sigma Cat# CS0030) according to the manufacturer’s instructions. Briefly, post-treatment cells were washed twice with PBS and fixed for 10–15 min with 10x fixation Buffer at room temperature. After fixation, cells were incubated with X-gal solution containing staining mixture at 37°C overnight in a dry incubator (without CO_2_). β-gal–positive cells were analyzed using a phase contrast microscope (OLYMPUS IX73). Results are plotted using mean ± SD of three independent experiments.

### Annexin V Analysis

Apoptosis was assessed using Annexin V staining according to the manufacturer’s instructions (ThermoFisher, Cat #A13199). Briefly, cells were seeded at a density of 2.5 x 10^5^ cells/mL in 6- well plates. The following day, the cells were treated with VEH, TAM (10 µM), DOXO (0.1 µM), and the combination of both for 24 h. Cells were harvested, washed 2x with ice cold phosphate-buffered saline, washed 1x with annexin V binding buffer and resuspended in 1x annexin v binding buffer at a concentration of 1 × 10⁶ cells/mL. A total of 100 µL of the cell suspension was incubated with FITC conjugated (Invitrogen, Cat #A13199) in the dark for 15 minutes. After incubation, 400 µL of 1× binding buffer containing 0.01 µg DAPI (ThermoFisher, Cat #62248) was added, and the samples were immediately analyzed by flow cytometry. At least 50,000 events were collected per sample using BD LSR Fortessa flow cytometer. Data were analyzed in FCS Express 7, and the cells were classified as: necrotic cells (Annexin V^-^/DAP^+^), late apoptotic cells (Annexin V^+^/DAP^+^), alive cells (Annexin V^-^/DAP^-^), and early apoptotic cells (Annexin V^+^/DAP^-^). The results are presented as mean values ± standard deviation, based on three independent experiments.

### Analysis of RNA-seq data

Sequencing reads that passed the quality filter from Illumina RTA were initially processed using FASTQC (v0.10.1) for base quality control. The sample reads were then aligned to the human reference genome (GRCh38) and the GENCODE (version 38) annotation database using STAR2(99) A second round of quality control was performed using RSeQC (100) on the mapped BAM files to identify any potential issues with the RNASeq library preparation. Gene-level raw counts were generated using the Subread package (101). Differential gene expression analysis was conducted using DESeq2 (102), and pathway analysis was carried out using the Gene Set Enrichment Analysis (GSEA) method (version 4.0) (103). The GSEA tool was selected to analyze the normalized gene count data, with low-count genes pre-filtered. Pathway analysis was performed against MSigDB, a curated collection of annotated gene sets provided by the developers of GSEA (Broad Institute, MIT and Harvard). The analysis utilized the canonical pathway collection from the version 2024 set.

### Immunofluorescence (IF) assay

Cells were seeded on sterile glass coverslips at a density of 2 × 10⁵ cells/well in 6-well plates. Following treatment, cells were washed with PBS and fixed in 4% paraformaldehyde (PBS, pH 7.4) for 15 min at room temperature. After washing, cells were permeabilized with 0.5% Triton X- 100 for 10 min and blocked with 3% non-fat milk for 1 h. Coverslips were incubated with primary antibodies (1% milk-PBS) for 3.5 h, washed 3x with PBS, and then incubated with secondary antibodies (0.5% milk-PBS) for 1.5 h. Nuclei were counterstained with 1 µg/mL DAPI for 15 min. After final washes, coverslips were mounted using ProLong™ Gold antifade medium (Invitrogen, Cat# P36930). Details of Primary and secondary antibodies and their dilution used for IF are provided in Table S1.

### Immunohistochemistry (IHC) analysis

Tissue microarray (TMA) was generated using 1-mm tissue cores (in triplicate) from formalin-fixed paraffin-embedded (FFPE) block of MDA231/LM2-4 CDX tissues. The antigen retrieval process was as previously described (98). High pH and low pH buffers were used for p53 and Ki67 staining, with low pH buffer applied for ERβ. The slides were incubated with Ki67 (R&D Systems, Cat# MAB7617) and p53 (Santa Cruz Biotechnology, Cat# sc-126) antibodies at a 1:50 dilution for 30 minutes, followed by treatment with Flex HRP polymer (Dako, Cat# DM843) for 30 minutes. For ERβ staining, the ERβ antibody (GeneTex, Cat# GTX83627) was applied at a 1:200 dilution for 20 minutes, followed by incubation with Flex HRP polymer for 20 minutes. Details of the antibodies used are provided in Table S2. Following immunohistochemical staining, the TMA slides were dehydrated through graded alcohols, cleared in xylene, and coverslipped using a permanent mounting medium. After the mounting medium had dried completely, the slides were examined to ensure that the tissue sections were intact and suitable for scanning. The mounted TMA slides were then scanned using the Leica Aperio ScanScope XT whole-slide imaging system. (ImageScope version 12.4.3.7009, Aperio Technologies, Inc., Vista, CA, USA) and scored based on the methodology outlined previously (98). The H-score was analyzed to measure the expression levels of the target proteins, and the results were plotted using GraphPad Prism version 9.

### Proximity Ligation Assay (PLA)

Fluorescence-Based PLA: In situ proximity ligation assay (PLA) in cultured cells was performed using the Duolink II reagent kit (Millipore-Sigma, St. Louis, MO, USA, Cat# DUO92008). Treated or untreated cells were fixed with freshly prepared 2% paraformaldehyde (Millipore-Sigma, St. Louis, MO, USA, Cat# P6148), blocked with 2% BSA, and permeabilized with 0.1% Triton-X-100. Cells were incubated with ERβ, p53, and p73 antibodies for 1 hour at room temperature. After primary antibody incubation, secondary probe hybridization, ligation, and amplification were performed according to the manufacturer’s instructions. Coverslips were mounted using ProLong Gold Antifade (Thermofisher Scientific #Cat P36930). Images were captured using an OLYMPUS IX73 microscope. PLA signals were quantified 10 different microscopic fields of three independent experiments and statistically analyzed by plotting using GraphPad Prism 9.

Bright-Field PLA: FFPE tissue TMAs (CDX and PDX) were deparaffinized and processed for bright-field PLA assays using Duolink® In Situ Detection Reagents Brightfield (Sigma-Aldrich, Cat# DUO92012), as previously described (91). Relevant pairs of primary antibodies against ERβ, p53, and p73 (dilution-1:100) applied to each corresponding TMA and incubated for 1 hour in a humidified chamber. After primary antibody incubation, mouse PLUS and rabbit MINUS secondary PLA antibodies were diluted and applied according to the manufacturer’s instructions. Following PLA staining, the TMA slides were dehydrated through graded alcohols, cleared in xylene, and coverslipped using a permanent mounting medium. After the mounting medium had dried completely, the slides were examined to ensure that the tissue sections were intact and suitable for scanning. The mounted TMA slides were then scanned using the Leica Aperio ScanScope XT whole-slide imaging system. ERβ–mutant p53 and mutant p53–p73 interaction signals were quantified as the number of PLA dots per 300 nuclei in tumor area per tissue core, across three independent experiments. Details of the antibodies and their respective dilutions are provided in Table S2.

### Lipid Peroxidation Assay

MDA231/LM2-4^LUC+^ and MDA-MB-468 cells were seeded at a density of (0.5 × 10^6^ cells/well) in a 6 cm dish. The following day, cells were treated with VEH, RSL3 (0.5 μM), Erastin (0.25 μM), TAM (10 μM), DOXO (100 nM), and the combination of both TAM and DOXO for 24 h. Post treatment, cells were washed 3x with PBS and incubated BODIPY 581/591 C11 (2 µM) in a CO_2_ incubator at 37°C for 1 hour. Cells were washed twice with cold PBS, resuspended in PBS containing 2% FBS and immediately analyzing using BD LSRFortessa flow cytometer. BODIPY 581/591 C11oxidation was assessed by measuring the shift from red fluorescence (non-oxidized: PE channel) and (oxidized: FITC channel). A minimum of 20,000 events per treatment were acquired and the data were analyzed using FCS Express 7. The lipid peroxidation was quantified as the ratio of oxidized (FITC)/ non-oxidized (PE) and plotted using GraphPad Prism 9. Microscopic images were acquired using OLYMPUS IX73. Stained cells are shown (Oxidized lipid: green; non-oxidized lipids: red, scale bar,100μm).

### Cell Viability Assay and IC_50_ Measurement

Half-maximum inhibitory concentration (IC_50_) values were measured using resazurin (Sigma Aldrich, Cat# R7017) assay. Briefly, Cells were seeded in a 96-well plate at a density of 10,000 cells/well. The following day, cells were treated with serial dilutions of DOXO ranging from 0 to 16 µM without and with TAM (10 µM) for 48 h. After treatment, 0.2 mM resazurin (Sigma Aldrich, Cat# R7017) was added, and the cells were incubated for 4 h at 37°C in a humidified incubator with 5% CO₂. Fluorescence was measured using BioTek Synergy H1 microplate reader with excitation at 540 nm and emission at 590 nm. The cell viability of the negative control (VEH-treated cells) was set at 100%, and the percent cell viability was calculated using the formula: [(experimental absorbance – background absorbance)/(absorbance of untreated control – background absorbance)] × 100%. Dose–response curves were constructed using nonlinear regression analysis in GraphPad Prism 9 (log[inhibitor] vs. response, variable slope). Experiments were performed in triplicate.

### Nuclear and Cytoplasmic Fractionation

After treatment, cells were harvested in cold cytosolic extraction buffer, vortexed, and centrifuged at 14,000 rpm for 5 minutes. The supernatant was collected as the cytosolic fraction, while the pellet was resuspended and vortexed with nuclear extraction buffer, followed by centrifugation at 14,000 rpm for 5 minutes. The nuclear extract was transferred and filtered into a pre-chilled collecting tube provided with the kit (Invent Biotech, Cat.# SM-005). The protein expression levels of ERβ, p53, Lamin A, and GAPDH were analyzed by immunoblotting.

### In-Vivo Mouse Modeling

All studies were approved and conducted in accordance with the Roswell Park Institutional Animal Care and Use Committee (IACUC) regulations (Protocol-1407M) and the National Institutes of Health (NIH) Guide for the Care and Use of Laboratory Animals. Mice were housed under pathogen-free conditions in an American Association for the Accreditation of Laboratory Animal Care (AAALAC)-accredited facility at Roswell Park.

Cell Line-Derived Xenograft (CDX) models: For orthotopic tumor implantation, MDA231/LM2- 4^LUC+^, MDA231/LM2-4^LUC+^ p53KO, MDA231/LM2-4^LUC+^ exogenous p53R248Q, and MDA231/LM2-4^LUC+^ p53KO exogenous wild-type p53 (0.75 × 10⁶ cells per mouse) were orthotopically implanted with a PBS: Matrigel mixture (Cat# 356234, Corning®, NY, USA) (1:1) into the 4th mammary fat pad of 6 week-old female SCID mice. Mice were randomized once the tumors reached approximately 200 mm³, and treatment was initiated. Mice were treated with tamoxifen (80 mg/kg) five days a week and doxorubicin (1 mg/kg) once a week, with vehicle controls of corn oil for tamoxifen and phosphate-buffered saline (PBS) for doxorubicin. Tamoxifen was administered by oral gavage five days a week, and doxorubicin was administered by intraperitoneal injection once a week. Tumor volumes were measured with calipers and calculated using the equation (width² × length)/2. For treatment preparation, doxorubicin was dissolved in water, and tamoxifen was dissolved in corn oil (Cat# C8267, Millipore Sigma, USA). All CDX study treatments were discontinued once the vehicle group tumors reached 2000 mm³, after which necropsy was performed. Following luciferin injection (10 mg/ml D-Luciferin, Cat# LUCK-100, Gold Biotechnology, St. Louis, MO, USA), metastasis analysis in the lung, liver, kidney, and brain was performed using IVIS imaging. Tumor tissues and collected organs were preserved for downstream analysis.

Patient Derived xenograft (PDX) models: All patient-derived xenograft (PDX) animal studies were conducted Baylor College of Medicine in accordance with the standard procedures approved by the Baylor College of Medicine Institutional Animal Care and Use Committee. For mammary gland

transplantation of PDX tumors, TNBC BCM3204, 3107, FCP699, 4849, 4013, 4913, and weak ER-positive TNBC BCM7441 were transplanted into the fourth mammary fat pads of 4-5-week-old SCID/Beige female mice. For the treatment study, doxorubicin was dissolved in water, and tamoxifen was dissolved in corn oil. Once the PDX tumors reached a size of 100-150 mm³, the mice were treated with tamoxifen (80 mg/kg) via oral gavage or doxorubicin (1mg/kg) via intraperitoneal injection. Tamoxifen treatment was administered 5 days a week, while doxorubicin treatment was given every two weeks. Tumor volumes were measured using calipers.

### PK/PD Model Development

Mean pharmacokinetic (PK) data for tamoxifen in mice following either a single oral or i.v. dose was obtained from the literature (104–106). Mean plasma doxorubicin concentrations in mice following either a single i.v. or i.p. dose were also extracted (107,108). Concentration-time data for both drugs were within a range expected based on the doses selected for our *in vivo* experiments, and the PK data were digitized using WebPlotDigitizer (https://automeris.io/WebPlotDigitizer/). A standard linear two-compartment model with first-order absorption and elimination processes was used to characterize the PK for both drugs.

CDX PD Model Development: A sequential PK/PD modeling approach was applied in which PK data were first modeled and then associated PK parameters were fixed in modeling the CDX pharmacodynamics. For tumor growth kinetics, a stepwise modeling approach was used, in which tumor growth and drugs as single agents were modeled first, and parameters associated with individual drug effects and tumor growth were fixed. For the drug combination, a modified form of the non-competitive interaction equation was used to determine the nature and extent of the interaction with a single parameter (*ΨCOME*) (109).

An exponential growth model was used to describe tumor growth in the absence of drug, where tumor volume (TV) proliferates via a first-order rate constant for growth (k_g_):

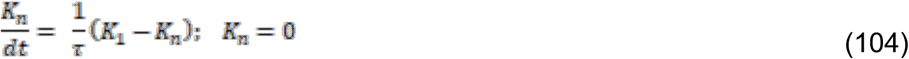

with TV_0_ representing the initial tumor volume size at time of treatment. Drug effects on tumor growth were characterized by a delayed cell kill signal (K_n_), which is driven by a series of transit compartments to capture the delay between drug exposure and effect: (110)

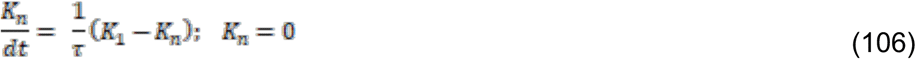

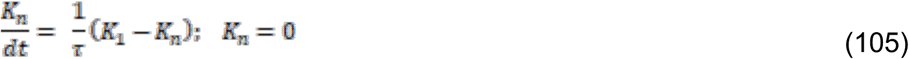

where C is drug concentration, k_kill_ is a linear proportionality constant, K_1_ and K_n_ represent a series of transit compartments to delay drug effect, with K_n_ being the final transit compartment (i.e., K_4_ in a four-transit compartment delay), and τ represents the mean transit time between compartments.

For the drug combination, a modified form of the non-competitive interaction equation was used to determine the nature and extent of the interaction with a single parameter (*ΨCOME*), in which the addition of the second drug impacts the potency of the first drug (109). The single agent drug model was modified to include the interaction parameter:

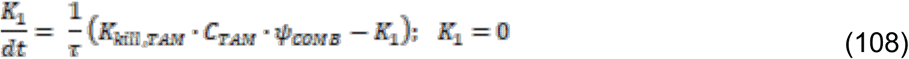

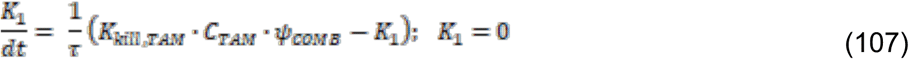

Based on this parameterization, an estimate of *ΨCOME* > 1 suggests synergism, *ΨCOME* < 1 suggests antagonism, and *ΨCOME* = 1 suggests an additive interaction. The interaction parameter was tested on each drug to determine if the nature of the interaction was the same. It was assumed that doxorubicin and tamoxifen operate via independent mechanisms of action. The final model allowed for a semi-mechanistic approach to determining the nature of the drug interaction. For PK and CDX data, model fitting was done using a naïve pooled approach, and parameter estimation was conducted using the maximum likelihood estimation method in ADAPT 5 [17]. A combinational variance model was utilized defined as: VAR = (σ_1_ + σ_2_•Y)^2^ where σ_i_ represents the estimated variance parameters, and Y is the model predicted value for drug concentration or tumor volume. Model evaluation and selection for PK and CDX analyses were guided by model fitting criteria such as visual inspection of model fitting, examination of goodness of fit plots, precision of parameter estimates, and the Akaike Information Criterion (AIC).

PDX PD Model Development: Owing to the high degree of variability in the PDX data, nonlinear mixed-effects (NLME) modeling was used to characterize tumor growth, treatment response, and the quantification of variability between individuals and treatment occasions. Interindividual variability (IIV) was tested for each parameter, and inter-occasional variability (IOV) was tested as needed. Both were assumed to follow a log-normal distribution:

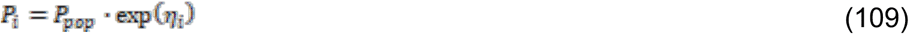

where *p_i_*represents the individual parameter estimate for individual *i*, *p_pop_* represents the typical population parameter estimate, and *η_i_* represents IIV or IOV random effects for the *i*th individual, which is normally distributed with a mean of 0 and a variance ω^2^. Occasions were defined as between doxorubicin doses. IIV was applied to all model parameters, except for *k_kill Dox_*, owing to large variability in the decline of tumor size following dosing events, and IOV was applied following each dose.

One categorical covariate of PDX ID was assessed using stepwise forward inclusion and backward elimination procedure to determine if covariate effects could be included on model parameters Covariates were included if, during forward selection, it reduced the objective function value (OFV) by at least 3.84 units and increased OFV by 6.64 units during backward elimination. The covariate model described as:

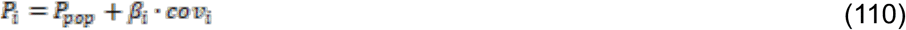

where *p_i_* represents the model-predicted parameter value for the *i*th individual, *p_pop_* represents the typical parameter value across the population, β*_i_* is the covariate effect, and *cov_i_* is an indicator variable that is fixed to 1 for covariate groups or fixed to 0 if covariate effects are not included.

Untreated tumor growth was described using a function, *g*(*t*), that accounts for two phases of tumor growth (Eq. in Supplemental Fig. S7A-E); an initial exponential growth phase, followed by a transition to a linear growth phase. A correlation was established between the linear tumor growth rate and initial tumor size and was included in the model. Single-agent drug effects for tamoxifen and doxorubicin were described using the same pharmacodynamic model as described previously (*K_i_* terms in Eqs. 1-3) with minor modifications. Doxorubicin therapy was best fit with three-transit compartments to account for the delay between concentration and effect, and no transit compartments were needed for tamoxifen. The combination model was described as:

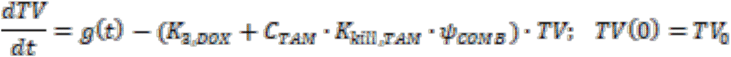

NLME modeling was performed using Monolix (version 2021R1) with the stochastic approximation expectation maximization (SAEM) algorithm as the estimation method. Residual unexplained variability was chosen among additive, proportional, and combined additive plus proportional models. Model selection and evaluation for PDX analysis was based on model fitting criteria, such as visual inspection of model fitting, goodness of fit plots, visual predictive check plots, precision of parameter estimates, and change in objective function. All data was plotted using the GraphPad Prism 8.0 software.

### Statistical Analysis

Data are presented as mean ± standard deviation (SD) from three biological replicates. Quantification of data was performed using GraphPad Prism version 9, Image J and FCS Express 7, as appropriate. Comparisons between two groups were performed using a two-tailed unpaired Student’s *t*-test, whereas comparisons involving two independent variables were performed using two-way analysis of variance (ANOVA), as appropriate. A *P* value ≤ 0.05 was considered statistically significant. Statistical significance is indicated as follows: \**P* ≤ 0.05, **P ≤ 0.01, \*\*\**P* ≤ 0.001, and ***P ≤ 0.0001. Representative immunoblotting and immunohistochemistry (IHC) images are shown in the corresponding figures. Each experiment was independently repeated at least two or three times, as indicated in the figure legends.

## Data Availability

All data supporting the findings of this study are included within the article and its Supplementary Information files. The RNA-sequencing (RNA-seq) datasets generated during this study are currently being deposited in a public repository, and the corresponding accession number will be provided upon availability. Additional data or materials required to reproduce or further interrogate the results will be made available by the corresponding author upon reasonable request.

## Authors’ Disclosures

No Disclosures.

## Acknowledgments

The work was supported by grants National Cancer Institute (NCI) at the National Institutes of Health (NIH) (R21CA137635, R01CA079911, and R01CA251545 to GMD), DOD BCRP Breakthrough Award (BC180932 to GMD and BAK), Roswell Park Alliance Foundation Grant to GMD, and NCI grant P30CA016056 involving the use of Roswell Park Comprehensive Cancer Center’s Shared Resources. We thank Drs. Michel Lewis and Lacey E. Dobrolecki, and the Baylor College of Medicine Patient-Derived Xenograft (PDX) Core for providing the PDX models. The PDX core is supported by the Cancer Prevention and Research Institute of Texas (CPRIT) Core Facility Award (RP220646) and the P30 Cancer Center Support Grant (NCI-CA125123).

## Notes

### Competing Interest Statement

The authors have declared no competing interest.

