## Supplemental data for "Tamoxifen and Doxorubicin Synergistically Target Estrogen Receptor β-Mutant p53-p73 Axis, a Therapeutically Actionable Vulnerability in Triple Negative Breast Cancer"

### **Supplemental Figures**

#### Supplemental Figure S1.

**Related to main Fig. 1; Tamoxifen stabilizes ERβ and enhances its interaction to Mutant p53 in TNBC cells.**

**A**, MDA-231/LM2-4 cells were transiently transfected with non-specific control siRNA (si-NS) or ERβ specific siRNA (siERβ) and empty vector (EV) and ERβ overexpression vector (ERβOV) for 48 h. Post transfections, cells were lysed by RIPA buffer and ERβ and β-actin expression level were analyzed by immunoblotting using ERβ specific PPZ0506 antibody (ThermoFisher Scientific Cat#MA5-24807). **B&C**, MDA-231/LM2-4 cells were treated with Veh and differential concentrations (25, 50, 75 nM) of PHTPP (**B**) and DPN (**C**) for 24 h. Immunoblot shows the expression of p53 and β-actin as a loading control. Right panels: Quantification of p53 protein expressions post PHTPP (**B**) and DPN (**C**) treatment analyzed by GraphPad Prism 9. **D**, MDA-231/LM2-4 cells were treated with Veh, Estradiol (50 nM) for 0, 12, and 24 hrs. Post treatment, cells were fixed by 10% formalin and subjected to proximity ligation assay (PLA) using ERβ- and p53-specific antibodies. Representative images display ERβ-mutant p53 interaction as pink fluorescent dots. Scale bar= 10 μm. Right panel: Quantification of average PLA signal (pink dots) per cell, counted manually and plotted using GraphPad Prism 9. **E**, Representative images showing the absence of PLA signal in the single-antibody control using anti-p53 (DO-1) alone. **F**, MDA-231/LM2-4 cells were treated with VEH and 50 nM DPN and Estradiol (E2) for 0, 12 & 24 hrs. Post treatment, cells were lysed using RIPA buffer, and the protein expression levels of ERβ and β-actin were assessed by immunoblotting. **B&C**, represents the mean ± SD from three independent experiments. Statistical significance was calculated using one-way ANOVA (ns=non-significant). Immunoblots in (**F**) are from one representative experiment with DPN and E2 treatment.

### Supplemental Figure S1

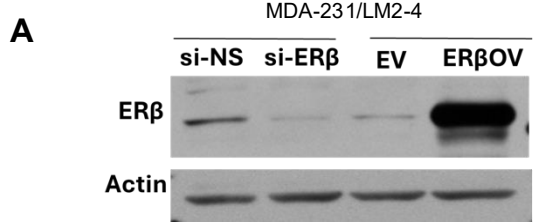

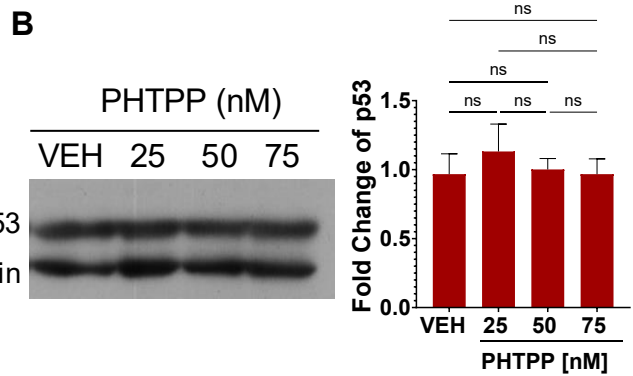

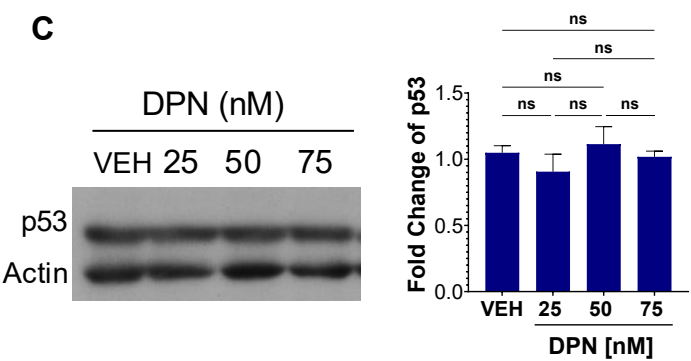

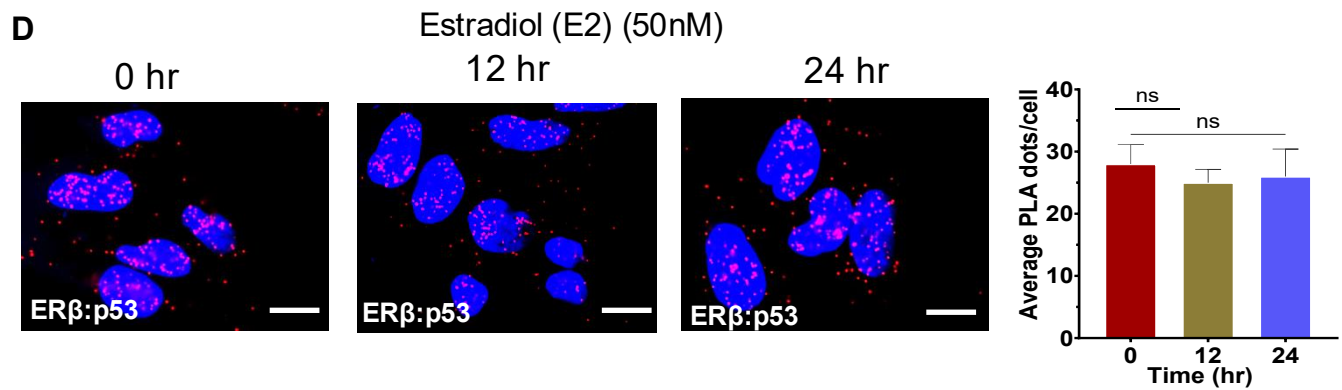

#### **E** <u>PLA: Single antibody control Control</u>

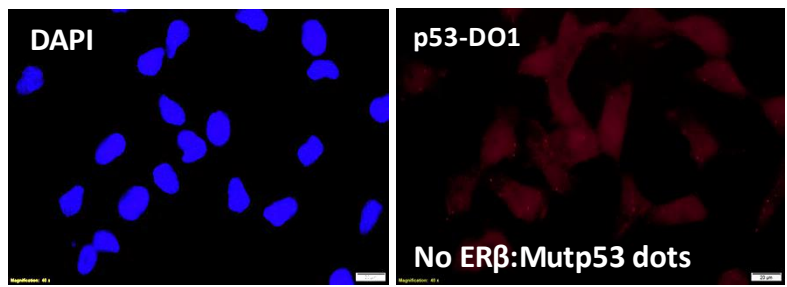

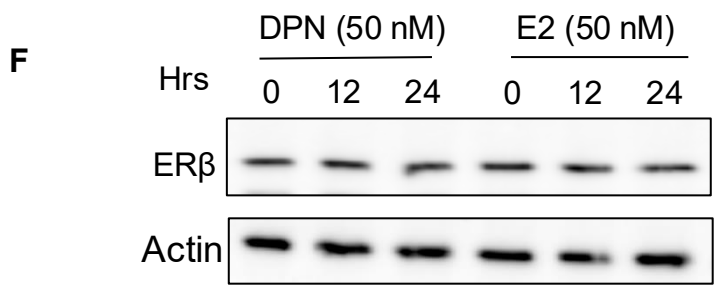

#### Supplemental Figure S2.

##### Related to main Fig 2; Mutant p53 Is Necessary for anti-tumorigenic Effects of Tamoxifen in TNBC cells.

**A&B**, MDA-231/LM2-4 (**A**) and MDA-231/LM2-4 p53 knockout (p53KO) (**B**) cells treated with Veh, or the ERβ-selective agonist diarylpropionitrile (DPN; 50 nM) for 24 h. Relative mRNA expression levels of p53 target genes *CDKN1A* (p21) and *BBC3* (PUMA) were determined by qRT-PCR and normalized to the housekeeping gene. **C&D**, MDA-231/LM2-4(**C**) and p53KO (**D**) cells were treated with VEH, and 4-OHT (10 μM) for 24 hours. After treatment, cells were pulsed with 10 μM BrdU for 1 hour, stained with Alexa Fluor® 488 Mouse anti-BrdU antibody (BD Pharmingen™, Cat# 558599) and propidium iodide (PI), and analyzed by flow cytometry. Quantification was performed using FCS Express 7 software. **E&F**, MDA-231/LM2-4(**E**) and p53KO (**F**) cells were treated with VEH, 4-OHT (10 μM), DPN(50nM), PHTPP(50nM) for 24 hours. For apoptosis analysis, cells were stained with Annexin V and propidium iodide to identify early and late apoptotic cell populations. Representative flow cytometry plots and quantitative analyses of cell cycle distribution and Annexin V-positive cells are shown. Data acquisition was performed using flow cytometry, and quantification was conducted using FCS Express 7 software. **G&H**, MDA-MB-468 cells were treated as described in **E** and **F**. Following treatment, cells were harvested and processed for flow cytometric analysis of cell cycle progression (**G**) and apoptosis (**H**) as described in E&F. **I&J**, MDA-231/LM2-4(**I**) and p53KO (**J**) cells were treated with VEH, and PHTPP (50nM) for 24 hours. For cell cycle analysis, cells were fixed in 70% ethanol and stained with propidium iodide (PI) to determine DNA content and quantify the distribution of cells in the G0/G1, S, and G2/M phases of the cell cycle. Quantification analysis were performed as discussed in **E&F**. **K&L**, MDA-231/LM2-4(**I**) and p53KO (**J**) cells were treated with VEH, and DPN (50nM) for 24 hours. For cell cycle analysis, cells were fixed in 70% ethanol and stained with propidium iodide (PI) to determine DNA content and quantify the distribution of cells in the G0/G1, S, and G2/M phases of the cell cycle. Quantification analysis were performed as discussed in **E&F**. **M**, MDA-MB-468 cells were treated with Veh, 50 nM DPN and PHTPP for 24 hours. Post treatment cells were processed for cell cycle distribution analysis. Cells were fixed in 70% ethanol, stained with propidium iodide (PI), and % cell cycle distribution was as discussed in E&F.

Error bars represent the mean ± SD. For Fig. A, B, I, J, and K, statistical significance between two treatment groups was determined using an unpaired two-tailed Student's t-test. For Fig. M, statistical significance among multiple treatment groups was assessed using one-way ANOVA. (adjusted P values: \*P ≤ 0.05 significant, ns=non-significant).

### Supplemental Figure S2

**A** MDA-231/LM2-4

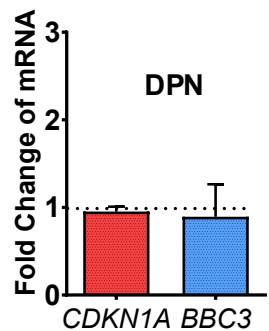

**B** MDA-231/LM2-4p53KO

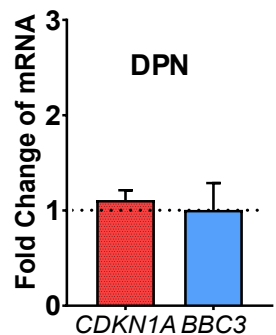

**C** Part of main Fig. 2E
MDA-231/LM2-4

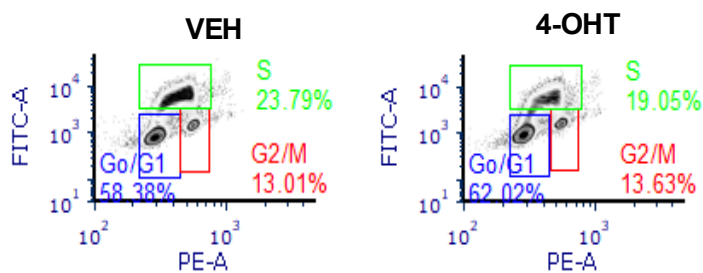

**D** Part of main Fig. 2G
MDA-231/LM2-4 p53KO

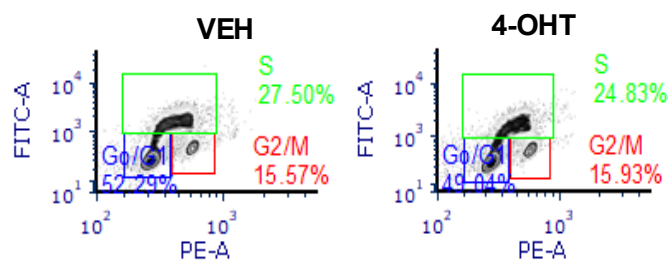

**E** Part of main Fig. 2F
MDA-231/LM2-4

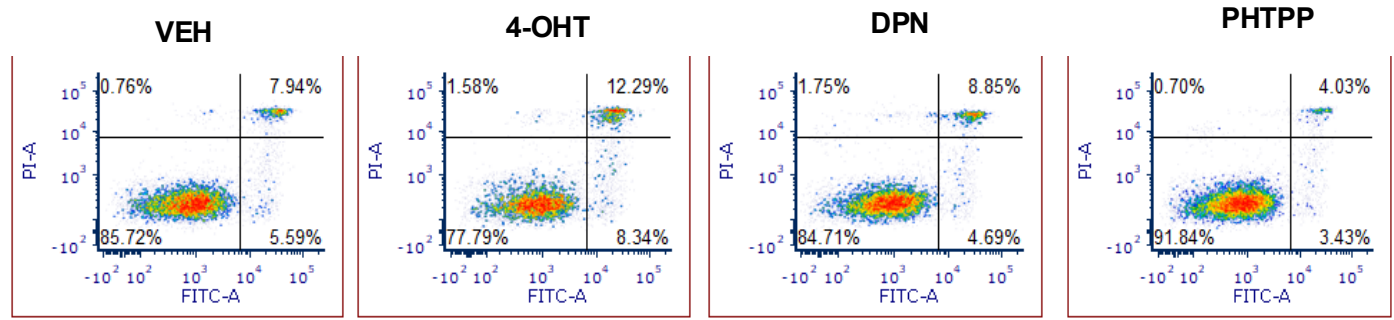

**F** Part of main Fig. 2H
MDA-231/LM2-4 p53KO

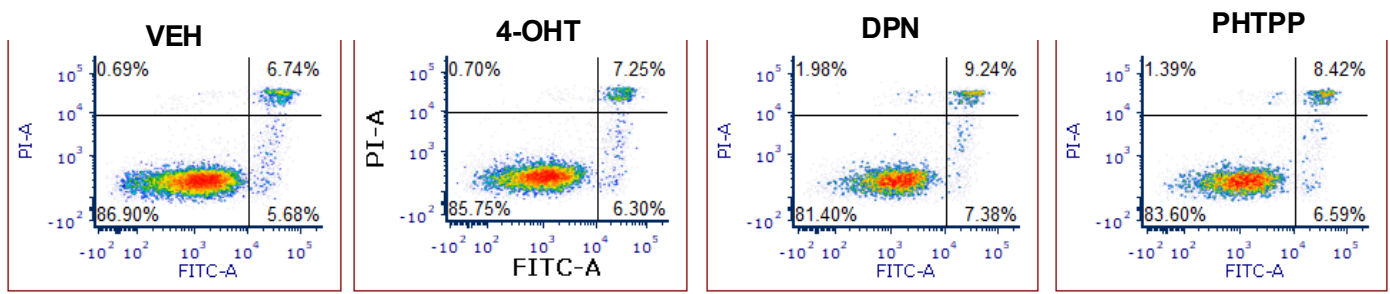

**G**

**Part of main Fig. 2J**

MDA-MB-468

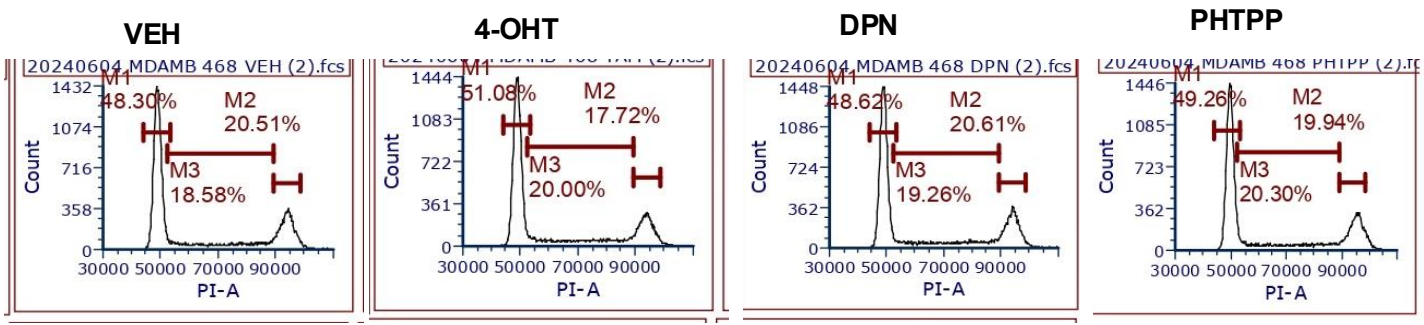

**H**

**Part of main Fig. 2K**

MDA-MB-468

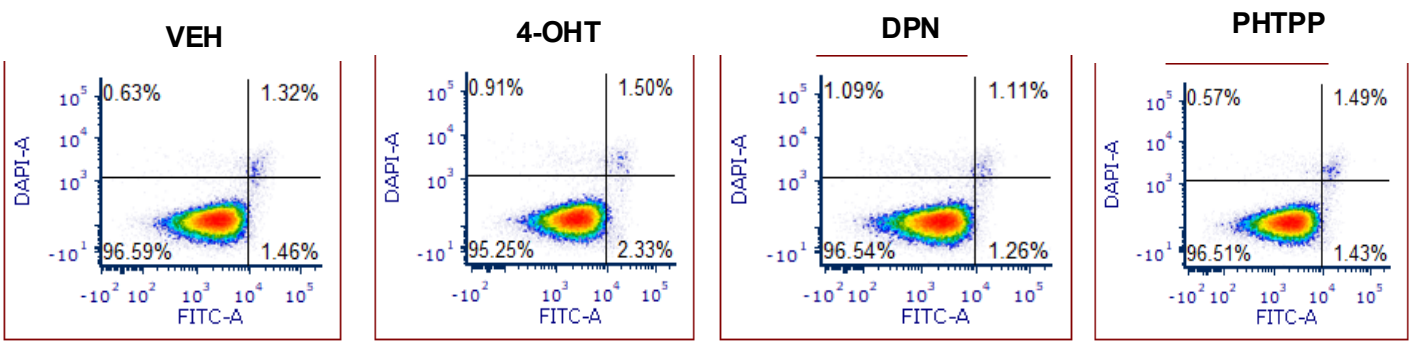

**I**

MDA-231/LM2-4

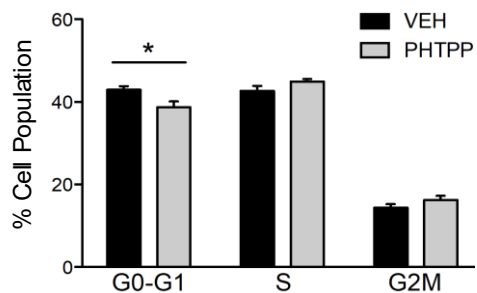

**J**

MDA-231/LM2-4 p53KO

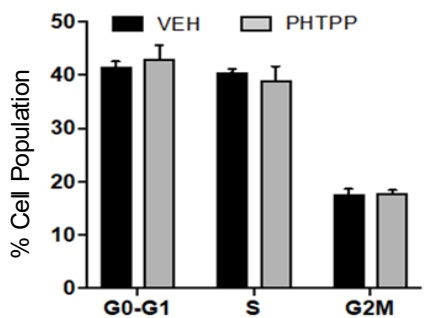

**K**

MDA-231/LM2-4

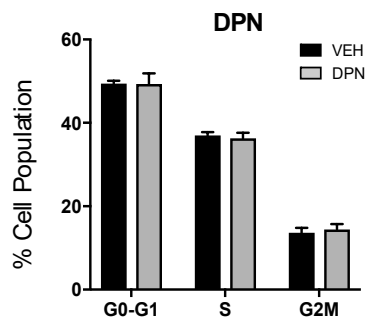

**L**

MDA-231/LM2-4 p53KO

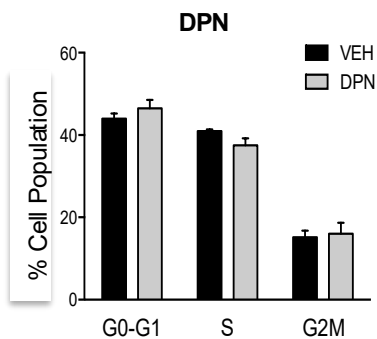

**M**

MDA-MB-468

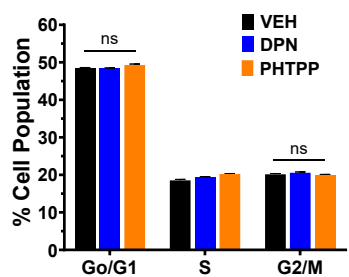

#### Supplemental Figure S3.

##### Related to main Fig. 3; Tamoxifen Synergizes with Doxorubicin to Induce Apoptosis and Suppresses Cell Proliferation, Migration, and Invasion in TNBC cells

**A**, Dose-response curves and IC<sub>50</sub> for DOXO and 4-OHT +DOXO showing growth inhibition of MDA-MB-468 cells. Cells treated with increasing concentrations of doxorubicin (DOXO) with Veh, 4-OHT (10 μM) for 48 h. **B&C**, Dose-response curves and IC<sub>50</sub> for 4-OHT showing growth inhibition of MDA-231/LM2-4 (**B**) and MDA-MB-468 (**C**) cells. treated with increasing concentrations of 4-OHT for 48 h. **D**, MDA-231/LM2-4 cells were treated with VEH, TAM (10 μM), DOXO (0.25 & 0.5 μM), or a combination of 4-OHT (10 μM) and DOXO (0.25 & 0.5 μM) for 48 hrs. Post 48hr, expression level of PARP, cleaved PARP, and actin were analyzed by immunoblotting. **E&F**, MDA-MB-468 (**E**) MDA-231/LM2-4(**F**) cells were treated with VEH, 4-OHT (10 μM), DOXO (0.5 μM), or combination of 4-OHT and DOXO for 48 hrs. Post-treatment, cell morphology was examined by phase-contrast microscopy. Representative images of surviving cells are shown. Scale bar =100 μm. **G &H**, MDA-231/LM2-4 p53KO cells following treatment with Veh, TAM (10 μM), DOXO (0.5 μM), or the combination of 4-OHT and DOXO for 24 hour. Post treatment, quantitative analysis of wound healing assay (**G**) from 0 to 24h matrigel invasion assay (**H**) from 0 to 20 h were analyzed and plotted using GraphPad prism 9. **I &J**, MDA-231/LM2-4 (**I**) and p53KO (**J**) cells were treated with VEH, 4-OHT (10 μM), DOXO (0.5 μM), or combination of 4-OHT and DOXO for 24h followed by senescence assay. Representative brightfield microscope images of senescence cells as shown (related to main Fig. 3 **O** and **P**). Scale bar = 50 μm.

Error bars indicate ± SD. Statistical analysis was performed using ANOVA in Fig S3 **H**. Data shown in **I** is set of a single experiment. (adjusted P values: \*\*\*P ≤ 0.0001, ns=non-significant).

Supplemental Figure S3

MDA-MB-468

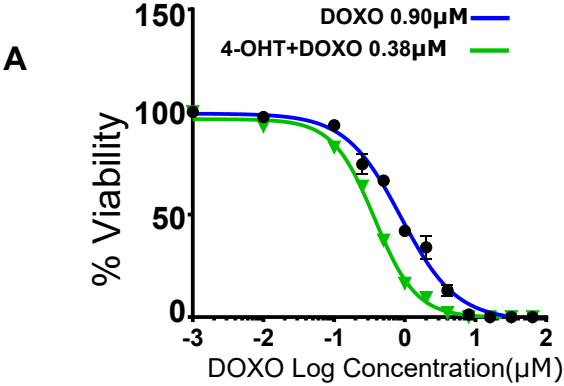

**B** MDA-231/LM2-4

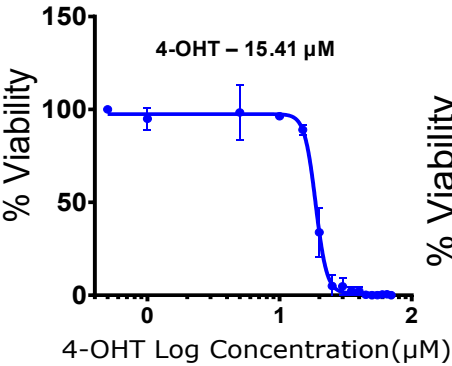

**C** MDA-MB-468

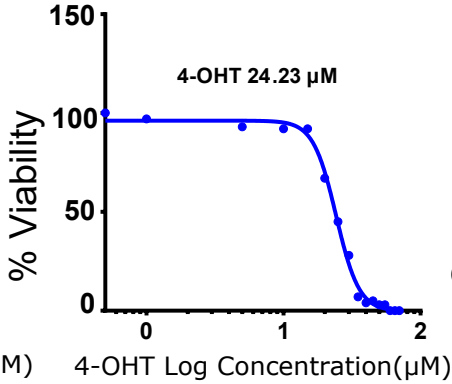

**D**

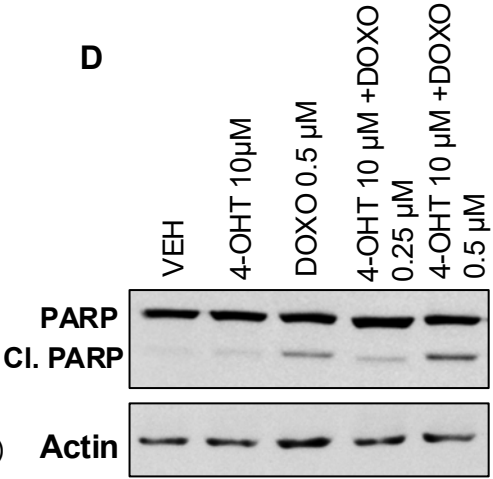

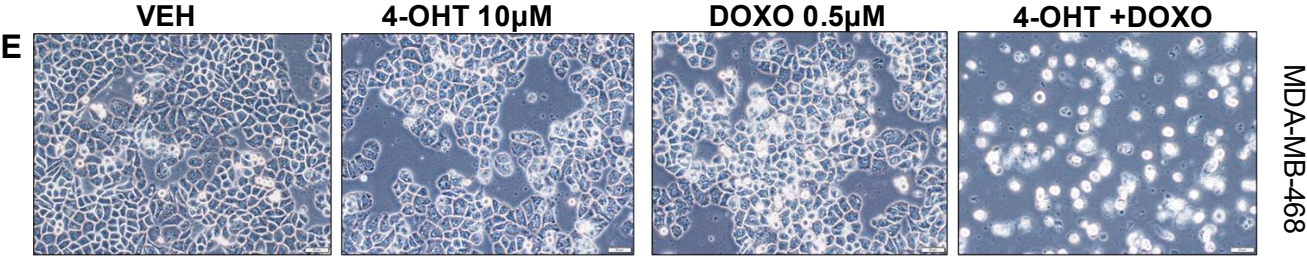

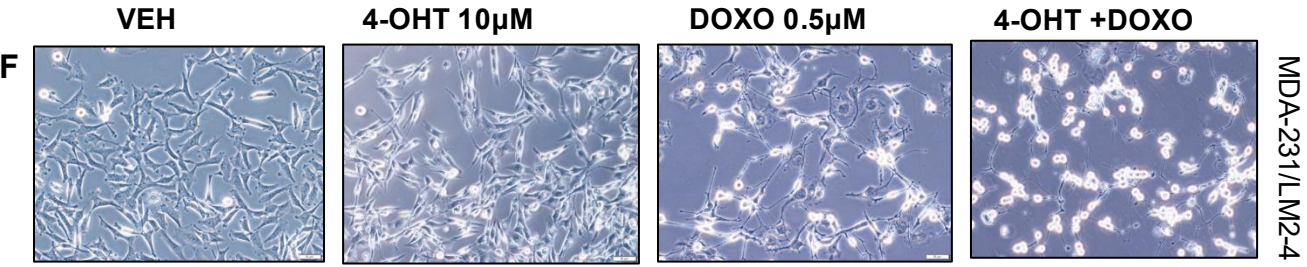

### Supplemental Figure S3 continued

MDA-231/LM2-4p53KO

Cell Migration Assay

G

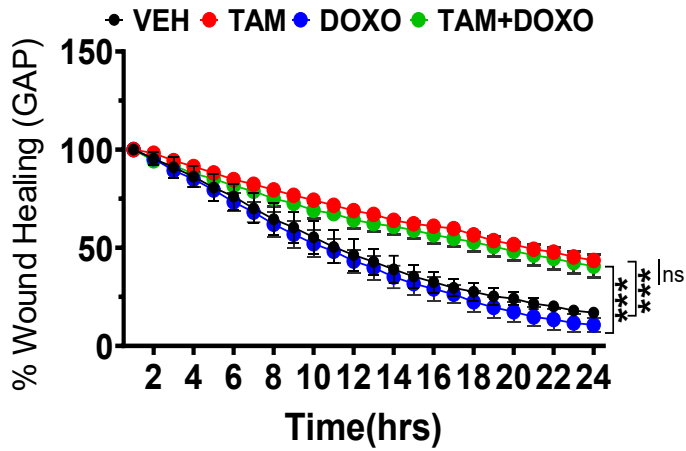

Cell Invasion Assay

H

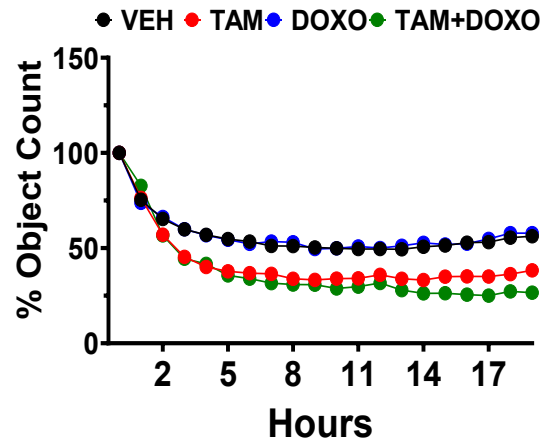

Senescence Assay: MDA-231/LM2-4

I

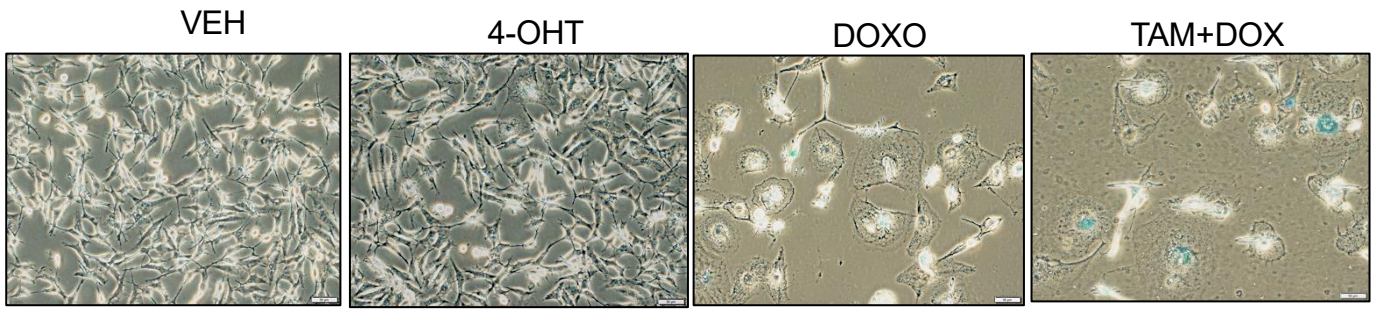

Scale Bar= 50 μm

Senescence Assay: MDA-231/LM2-4 p53KO

J

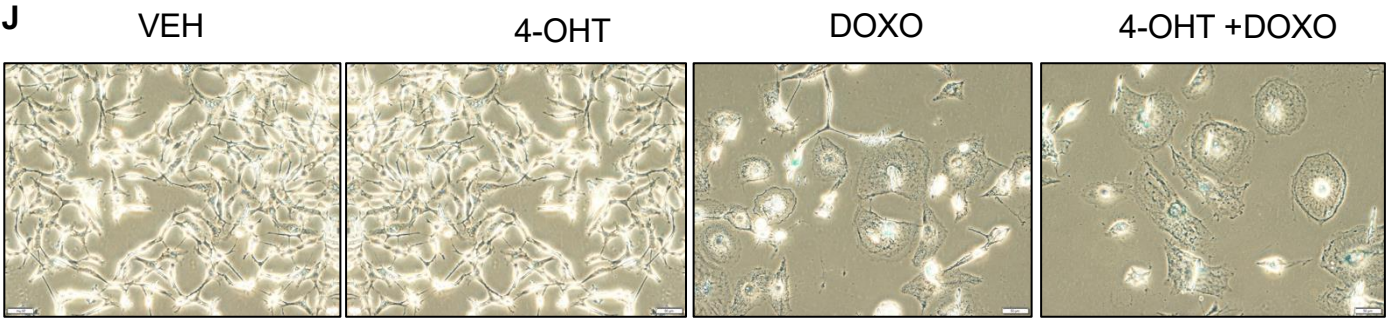

Scale Bar= 50 μm

#### Supplemental Figure S4.

##### Related to main Fig. 4; ERβ interaction with Gain-of-Function p53 Mutants Enhances Apoptotic Response to Combined Tamoxifen–Doxorubicin Treatment.

**A**, Schematic representation of the generation of isogenic MDA-231/LM2-4TNBC cell lines with different combination of ERβ, mutant and wild type p53. Right panel; MDA-231/LM2-4(endogenous Mutant p53R280K), MDA-231/LM2-4p53KO (p53-null; empty vector), and p53KO cells reconstituted with exogenous p53 mutants (R175H, R273H, R248Q, R280K) or wild-type p53 (WT) were cultured for 24 hours. Whole-cell lysates were prepared using RIPA buffer, and expression levels of ERβ, p53, and β-actin were analyzed by immunoblotting. **B**, ERβ binding ELISA assay: p53KO cells were transiently transfected with empty vector (control) and p53R175 and p53R248Q overexpression vector for 48 h. ERβ binding affinity was analyzed using ELISA assay. **C**, Dose response curves and IC<sub>50</sub> values of the following eight isogenic cells, p53KO, p53KO cells expressing p53WT, p53KO cells expressing different p53 mutants (R175H, R273H, R248Q, and R280K cells with ERβ knockdown), and p53KO cells with ERβ knockdown following treatment with increasing concentrations of doxorubicin (DOXO) and 4-OHT for 48h (Data analysis related to main fig 4.C). **D**, TNBC cells expressing p53 null (p53KO), wild-type p53 (p53WT), mutant p53R248Q, p53R273H, and p53R280K were treated VEH, 4-OHT (10 μM), DOXO (0.1 μM), or the 4-OHT + DOXO combination for 24 h, followed by Annexin V staining. Apoptotic (Annexin V–positive) cells were quantified by flow cytometry, and data were plotted using GraphPad Prism 9. (Data analysis related to main fig 4. **E**, MDA-MB-231/LM2-4 cells were transiently transfected with non-specific control siRNA (si-NS) or ERβ-specific siRNA (si-ERβ) for 48 h, followed by treatment with Veh, 4-OHT; 10 μM, DOXO; 0.1 μM, or the combination of 4-OHT (10 μM) and DOXO (0.1 μM). Apoptosis was assessed after 24 h of treatment using Annexin V staining, followed by flow cytometric quantification of Annexin V-positive cells. **F**, MDA-231/LM2-4 cells were treated with 4-OHT (10 μM), DOXO (0.5 μM), and combination for 24 hrs. Post-treatment, cells were fixed in 2% paraformaldehyde and permeabilized with 10% BSA + Triton-X100, followed by immunofluorescent (IF) assay using ERβ and p53 specific primary antibodies followed by secondary antibody Alexa Fluor 488 (Green). DAPI was used for nuclear staining. Representative IF images as shown in fig. Scale bar = 20 μm. **G**, p53R248Q cells were treated for VEH, 4-OHT(10 μM), DOXO (0.5 μM) and combination for 24hrs. Post 24 hr, cells pre-stained with live Mito-tracker green as per reagent protocol (Invitrogen, Cat# M7512) and fixed by 2% paraformaldehyde. Fixed cells were processed for ERβ and mutant p53 PLA. Representative images of ERβ and mutant p53 interaction (pink dots) merged with Mito-tracker (green) as shown in figure. Scale bar = 10 μm.

Supplementary Figure S4

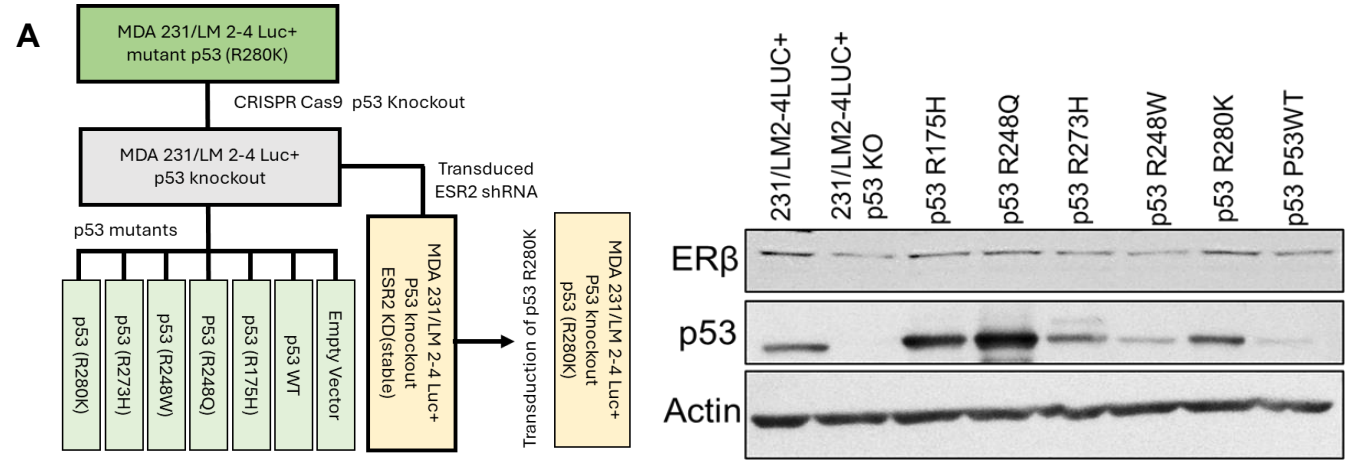

C

D

#### Part of main Fig. 4 C

MDA-231/LM2-4

p53KO

p53WTP53

p53R248Q

p53R273H

p53R280K

**E** MDA-231/LM2-4 Immunofluorescence

**F** MDA-231/LM2-4 p53R248Q

#### **Supplementary Figure S5.**

**Related to main Fig. 5; ERβ Sequesters Mutant p53 from p73 to Enhance Apoptotic Response to Tamoxifen–Doxorubicin Combination treatment.**

**A and B**, MDA-231/LM2-4 (**A**) and MDA-MB-468 (**B**) cells were transiently transfected with non-specific (si-NS) or p73-targeting siRNA (si-TP73) for 48 h, followed by treatment with VEH, 4-OHT (10 μM), DOXO (0.5 μM), or the 4-OHT + DOXO combination for 24 h. For apoptosis analysis, cells were stained with Annexin V and propidium iodide to identify early and late apoptotic cell populations. Representative flow cytometry plots and quantitative analyses of cell cycle distribution and Annexin V-positive cells are shown. Data acquisition was performed using flow cytometry, and quantification was conducted using FCS Express 7 software. (Flow analysis is related to main fig 5 **H and K**).

#### Part of main Fig. 5 H

MDA-231/LM2-4

#### Part of main Fig. 5 K

MDA-MB-468

#### Supplemental Figure S6

##### Related to main Fig. 6; Tamoxifen and Doxorubicin Combination Induces both Ferroptosis and Apoptosis

**A**, Venn diagram illustrating the overlap of upregulated and downregulated genes identified by comparative gene expression analysis of MDA-231/LM2-4 cells following treatment with Veh, 4-OHT, DOXO, or the combination of 4-OHT and DOXO for 24 hours.

**B-K** RNA sequencing analysis of MDA-231/LM2-4 cells treated with Veh, 4-OHT (10 μM), DOXO (0.5 μM), or the combination for 24 h (n=3, biological replicates). Representative GSEA plots

highlighting gene sets related to

HALLMARK\_E2F\_TARGETS (**B**),

HALLMARK\_MYC\_TARGETS (**C**),

REACTOME\_DNA\_REPAIR (**D**),

GOBP Mitochondrial Respiratory Chain Complex Assembly (**E**),

REACTOME regulation of cholesterol biosynthesis by SREBP (**F**), S

Ferroptosis Driver (**G**),

Ferroptosis Suppressor (**H**), REACTOME\_VOLTAGE\_GATED\_POTASSIUM\_CHANNELS (**I**),

REACTOME\_IRON\_UPTAKE\_AND\_TRANSPORT (**J**),

REACTOME Caspase activation via death receptor in the presence of ligand (**K**).

**L**, Heat map of mevalonate pathway associated genes generated from RNA-seq analysis, demonstrating their upregulation (with 4-OHT) and downregulation (with doxo and combination) following 24-hour treatment.

**M**, RNA-sequencing (RNA-seq) analysis of mevalonate pathway-associated gene expression in MDA-231/LM2-4 cells following the indicated treatments. Green arrows indicate downregulation, whereas red arrows indicate upregulation of gene expression.

**N&O**, Heat map data of Ferroptosis driver (**N**) and suppressor (**O**) in MDA-231/LM2-4 and generated from RNA-seq analysis of and following 24 h treatment with VEH, 4-OHT, DOXO and 4-OHT + DOXO.

**P&Q**, Selective ferroptosis driver (**P**) and Selective ferroptosis suppressor (**Q**) in MDA-231/LM2-4 generated from RNA-seq analysis of and following 24 h treatment with VEH, 4-OHT, DOXO and 4-OHT + Doxo.

**R&S**, MDA-231/LM2-4 and MDA-231/LM2-4 p53-knockout (p53KO) cells were treated with VEH, 4-OHT (10 μM), DOXO (0.5 μM), or a combination for 24 hours. Following treatment, *TFRC* mRNA (**R**) and TFR1 (**S**) protein expression was analyzed by qRT-PCR and immunoblotting, respectively.

**T & U**, MDA-231/LM2-4ERβ knockdown cells were treated with VEH, 4-OHT (10 μM), DOXO (0.5 μM), or a combination for 24 hours. Following treatment, *TFRC* (**T**) and *GPX4* (**U**) mRNA analyzed by qRT-PCR.

**V**, MDA-231/LM2-4 cells were treated with RSL3 (0.25 μM) for 24 hrs. Following treatment, cells were stained with BODIPY581/591 (2 μM) for 1 hr. Assessment of lipid peroxidation in cells using IF Assay. Stained cells are shown (Oxidized lipid: green; non-oxidized lipids: red, scale bar, 100 μm).

**W**, MDA-MB-468 cells were treated with VEH, RSL3 (0.25 μM), 4-OHT (10 μM), Doxo (0.5 μM), and TAM+ Doxo for 24 hrs. Following treatment, cells were stained with BODIPY581/591 (2 μM) for 1 hr. Assessment of lipid peroxidation in cells using IF Assay (I). Stained cells are shown (Oxidized lipid: green; non-oxidized lipids: red, scale bar, 100 μm).

Error bars represent ± SD. RNA-seq data were analyzed across three independent experiments. The qRT-PCR data shown in **R**, **T**, and **U** represent three independent experiments. Adjusted P values are indicated as follows: \*P ≤ 0.05, \*\*P ≤ 0.01, \*\*\*P ≤ 0.001; ns, not significant.

Supplemental Figure S6

MDA-231/LM2-4

4-OHT

DOXO

4-OHT +DOXO

**B**

HALLMARK\_E2F\_TARGETS

**C**

HALLMARK\_MYC\_TARGETS\_V1

REACTOME\_DNA\_REPAIR

**D**

4-OHT

DOXO

4-OHT+DOXO

E

#### GOBP Mitochondrial Respiratory Chain Complex Assembly

F

#### REACTOME\_regulation of cholesterol biosynthesis by SREBP\_SREBF

G

#### Ferroptosis Driver

H

#### Ferroptosis Suppressor

4-OHT

DOXO

4-OHT+DOXO

#### I REACTOME\_VOLTAGE\_GATED\_POTASSIUM\_CHANNELS

J

#### REACTOME\_IRON\_UPTAKE\_AND\_TRANSPORT

K

#### REACTOME Caspase activation via death receptor in the presence of ligand

**L**      Mevalonate Pathway related gene expression in MDA-231/LM2-4

**M**

Mevalonate Pathway in response to Tamoxifen and doxorubicin treatment
in MDA-231/LM2-4

**N** Expression of Ferroptosis Driver in MDA-231/LM2-4

Expression of Ferroptosis Driver in MDA-231/LM2-4

O

Expression of Ferroptosis suppressor in MDA-231/LM2-4

Expression of Ferroptosis suppressor in MDA-231/LM2-4

### P, Quantification of RNA-seq data showing the expression of selected ferroptosis driver genes.

#### TFRC

#### HMOX1

#### NCOA4

## TF

#### FADS1

#### ACSL5

#### LPCAT3

#### ALOXE3

#### POR

#### SOD2

### Q, Quantification of RNA-seq data showing the expression of selected ferroptosis suppressor genes.

#### SREBF2

#### ACAT2

#### PMVK

#### TRIT1

#### FDPS

#### SQLE

#### DHCR7

#### ACACA

#### NDUFS1

#### NDUFB8

#### NDUFV1

#### UQCRC1

#### UQCRC2

#### UQCRCF1

**R**

**S**

#### MDA-231/LM2-4ERβKD

**T**

**U**

#### Part of main Fig. 6 I

#### RSL3 treatment to MDA-231/LM2-4

**V**

**W**

#### Lipid peroxidation assay in MDA-MB-468

Table S1, Basal gene expression levels and changes in gene expression following combined treatment with 4-OHT and DOXO in Mutp53 cells.

| Genes | Basal level expression in Mutp53 context<br>as compared to null p53 context |  | Expression in response to treatment with 4-OHT +DOXO as compared to Vehicle |  | Drug response (4-OHT<br>+DOXO) dependency on<br>Mtp53 context |
| --- | --- | --- | --- | --- | --- |
|  | Change | Significance | Change | Significance |  |
| SREBF2 | No Change |  | DOWN | 0.0007 | IND |
| ACAT2 | No Change |  | DOWN | 0.0052 | IND |
| MVK | DOWN | 0.0052 | DOWN | 0.0013 | IND |
| MVD | No Change | No Change | No Change |  | DEP |
| FDPS | No Change |  | No Change |  | IND |
| TFRC | DOWN | 0.0007 | Up | 0.0055 | DEP |
| HMOX1 | DOWN | 0.04 | Up | 0.0005 | IND |
| FTH1 | DOWN | 0.0114 | Up | 0.0002 | IND |
| NCOA4 | DOWN | 0.0037 | Up | <0.0001 | DEP |
| TF | No Change |  | Up | 0.001 | DEP |
| ACACA | DOWN | 0.063 | DOWN | <0.0001 | IND |
| FADS1 | DOWN | 0.0043 | Up | 0.0011 | DEP |
| SCD | No Change |  | DOWN | 0.0002 | IND |
| ACSL5 | DOWN | 0.0043 | Up | 0.0012 | IND |
| LPCAT3 | DOWN | 0.0007 | Up | 0.0002 | DEP |
| ALOXE3 | UP | 0.0264 | Up | <0.0001 | DEP |
| POR | DOWN | 0.0073 | Up | <0.0001 | IND |
| SOD2 | No Change |  | Up | <0.0001 | IND |
| NDUFS1 | DOWN | 0.0178 | DOWN | <0.0001 | IND |
| NDUFB8 | DOWN | 0.026 | DOWN | 0.0013 | IND |
| NDUFS3 | DOWN | 0.0484 | No Change |  | DEP |
| NDUFV1 | No Change |  | No Change |  | IND |
| UQCRC1 | DOWN | 0.0194 | DOWN | 0.0488 | DEP |
| UQCRC2 | No Change |  | DOWN | 0.0053 | DEP |
| UQCRFS1 | DOWN | 0.0327 | DOWN | 0.0017 | DEP |
| E2F3 | DOWN | 0.0015 | No Change |  | DEP |
| IREB2 | DOWN | 0.0112 | Up | 0.0383 | DEP |
| TLR3 | DOWN | 0.004 | Up | 0.0437 | DEP |
| TRIM22 | No Change |  | No Change |  | DEP |
| HSPA5 | DOWN | 0.007 | DOWN | 0.0169 | IND |
| GPX4 | No Change |  | No Change |  | DEP |
| STC1 | UP | 0.0004 | DOWN | 0.0001 | IND |
| PIM1 | UP | 0.0006 | DOWN | 0.0008 | DEP |
| ASS1 | UP | 0.0004 | DOWN | 0.0042 | DEP |
| SLC2A12 | UP | 0.0001 | DOWN | 0.0017 | DEP |
| IGF1R | No Change |  | DOWN | 0.0022 | DEP |
| CXCL8 | DOWN | 0.0008 | DOWN | 0.0012 | DEP |
| SLC7A11 | UP | 0.0025 | Up | 0.0094 | IND |
| SLC3A2 | UP | 0.0024 | Up | <0.0001 | IND |
| CYCS | UP | 0.0135 | Up | <0.0001 | DEP |
| CFLAR | No Change |  | Up | <0.0001 | DEP |
| TNFRSF10B | No Change |  | Up | 0.0001 | DEP |
| TNFRSF10D | UP | 0.0431 | Up | 0.0002 | DEP |
| CASP8 | No Change |  | Up | 0.0003 | DEP |
| E2F7 | No Change |  | Up | 0.0006 | DEP |
| BCL2L11 | DOWN | 0.0001 | Up | <0.0001 | IND |
| BBC3 | No Change |  | No Change |  | IND |
| CDKN1A | DOWN | 0.0017 | Up | <0.0001 | IND |
| APAF1 | No Change |  | Up | 0.0337 | DEP |

#### Supplemental Figure S7

##### Related to main Fig. 7; Tamoxifen & Doxorubicin Combination Suppresses Tumor Growth in vivo.

**A**, Model-fitted pharmacokinetic (PK) profiles of tamoxifen and doxorubicin. Symbols represent digitized data from original publications; solid lines represent fitted model profiles. **B**, Schematic representation of the pharmacodynamic (PD) model showing the effect of tamoxifen and doxorubicin combination therapy. The model includes transit compartments leading to tumor volume decline. **C**, Goodness-of-fit assessment of the PK/PD model in immortal cell line combination data, comparing observed versus model-predicted values. **D**, Predictive distribution from the combination therapy model in the PDX study. The dashed lines indicate the 5th and 95th percentiles, the solid line indicates the median prediction, and individual data points are represented by symbols. **E**, Goodness-of-fit evaluation for the PDX model, comparing observed values with individual model-predicted values under combination therapy. **F**, Treatment schema for MDA-231/LM2-4 p53R248Q cell-derived xenograft (CDX) models: SCID mice were treated with vehicle (corn oil and PBS), tamoxifen citrate (TAM; 80 mg/kg, oral gavage), or doxorubicin (DOXO; 1 mg/kg, intraperitoneal injection). Tumor volumes were measured at the indicated time points. Data are presented as mean ± SEM (n = 5 mice per group). Tumor volume plot: Symbols (●, ■, ▲, ▼) represent observed tumor volume data, and lines represent the model-fitted values. **G**, Body weights of CDXs MDA-231/LM2-4, MDA-231/LM2-4 P53ko, MDA231/LM 2-4 ERβKD, MDA/231 LM 2-4 WTp53, MDA/231 LM 2-4 R248Q. **H**, Post termination of MDA-231/LM2-4CDX model, tumors tissue were utilized to generate tissue microarrays (TMAs). Immunohistochemistry (IHC) was performed on TMA to assess ERβ and p53 expression using specific primary antibodies. Representative IHC images of each treated conditions are shown. Scale bar = 100 μm. **I**, Immunoblotting analysis of ERβ, p53, and β-actin in PDX tumor tissues. **J**, Detailed information on patient-derived xenograft (PDX) datasets obtained from the PDX Portal, Baylor College of Medicine, Texas. **K**, PDXs with mutant p53 expression BCM7441 were treated with vehicle (corn oil and PBS), tamoxifen citrate (TAM; 80 mg/kg, oral gavage), or doxorubicin (DOXO; 1 mg/kg, intraperitoneal injection) and tumor volumes were measured at the indicated time points. **L** Body weights of PDXs BCM3204, BCM3107, BCMFCP699, BCM4849, BCM4013, and BCM4913. **M-O**, Immunohistochemical staining on BCMPDX3204 tissues ERβ (ThermoFisher PPZ0506, Cat# MA5-24807) (**M**), p53 (Santa Cruz, Cat# sc-126) (**N**), and Ki-67 (R&D Biosystems, Cat# AF7617) (**O**). Tumor volume data in panels A and K are presented as mean ± SEM. Statistical significance was determined using ANOVA. Adjusted P values are indicated as follows: \*\*P ≤ 0.001; ns, not significant.

Supplemental Figure S7

F MDA-231/LM2-4p53R248Q

Experimental schema

Tumor Volume

G

#### Body weight of CDX model mice during treatment

MDA-231/LM2-4

MDA-231/LM2-4p53KO

MDA-231/LM2-4 ERβKD

MDA-231/LM2-4WTp53

MDA-231/LM2-4p53R248Q

H

MDA-231/LM2-4 CDX model IHC

Supplementary Figure S7

**J**

| BCM-PDX | Age at Diagnosis (Years) | p53 Status | ERβ Expression | Race | Response to TAM+DOXO treatment |
| --- | --- | --- | --- | --- | --- |
| BCM3204 | 42.22 | Mutant p53/ I332_N345del | + | WHITE | RESPONDED |
| BCM3107 | 57.6 | Mutant p53/T155N | + | WHITE | RESPONDED |
| BCMFCP699 | 63.92 | Mutant p53/R249S | + | WHITE | RESPONDED |
| BCM7441 | 35.75 | Mutant p53/R248Q | + | WHITE | RESPONDED |
| BCM4849 | 59.57 | p53 Null | + | WHITE | NOT RESPONDED |
| BCM4013 | 46.93 | WTp53 | + | BLACK | NOT RESPONDED |
| BCM4913 | 37.59 | Mutant p53/R306* | + | BLACK | NOT RESPONDED |

**K**

L

Body weight of PDX model mice during treatment

BCM3204

BCM3107

BCMFCP699

BCM4849

BCM4013

BCM4913

BCM7441

### Supplementary Figure S7

#### PDX BCM3204- IHC

ERβ

M

VEH

TAM

DOXO

TAM+DOXO

Scale bar = 50 μm

p53

N

Scale bar = 50 μm

ki67

O

Scale bar = 100 μm

**Table S2 Antibody details**

| Biomarker | Antibodies | Dilutions |  |  |  |  | Company (Cat. No.) |
| --- | --- | --- | --- | --- | --- | --- | --- |
|  |  | IHC | PLA | IF | IB | IP |  |
| ERβ | 68-4 | - | - | - | (1:5000) | - | Millipore (05-824) |
|  | 14C8 | (1:300) | (1:200) | (1:200) | — | 5μg | GeneTex (GTX70174) |
|  | PPZ0506 | (1:100) | - | - | (1:5000) | - | ThermoFisher (MA5-24807) |
| TP53 | DO1 | (1:10000) | - | - | - | - | Santa Cruz (SC-126) |
|  | FL393 | - | (1:300) | (1:300) | - |  | Santa Cruz (SC-6243) |
| TP73 | p73 (D3G10) | - | - | - | (1:5000) | - | Cell Signaling (14620) |
| PUMA | Anti-PUMA | - | - | - | (1:5000) |  | Cell Signaling (12450) |
| β-actin | Anti-β-actin | - | - | - | (1:10000) |  | Sigma (A5441) |
| PARP | Anti-PARP | - | - | - | (1:5000) | — | Cell Signaling (9542) |
| p21 | p21 Waf1/Cip1 (12D1) | - | - | - | (1:5000) | — | Cell Signaling (2947) |
| BIM | Anti-BIM | - | - | - | (1:10000) | — | Cell Signaling (2933) |
| Caspase 3 | Anti-Caspase-3 | - |  |  | (1:10000) | — | Cell Signaling (9662) |
| Mitotracker | MitoTracker™ Green FM | - | - | (1:300) | - |  | Thermofisher (M7514) |
| HRP secondary antibodies | HRP-anti-rabbit | - | - | - | (1:10000) | - | Sigma Millipore (12-348) |
|  | HRP-anti-mouse | - |  |  | (1:10000) | - | Sigma Millipore (12-349) |
| IF secondary antibodies | Alexa Fluor 488 anti-mouse | - | - | (1:250) | - | — | Invitrogen (A11001) |
|  | Alexa Fluor 594 anti-rabbit | - | - | (1:250) | - | — | Invitrogen (A11012) |
|  | Anti-mouse IgG (H+L) | - | - | (1:250) |  |  | Cell Siganaling (4409S) |
| IHC secondary antibodies | HRP-conjugated anti-rabbit | (1:100) | - | - | (1:10000) | - | DAKO (K4003) |
|  | HRP-conjugated anti-mouse | (1:300) | - | - | (1:10000) | - | Leica (PV-6114) |
| IgG negative controls | Normal mouse IgG | - | - | - | - | 5μg | Millipore (12-371) |
|  | Normal rabbit IgG | - | - | - | - | 5μg | Millipore (12-370) |

IHC- Immunohistochemistry ,PLA-Proximity Ligation Assay, IF-Immunofluorescence, IB- Immunoblotting, IP-Immunoprecipitation
